# Incorporating topological and age uncertainty into event-based biogeography supports paleo-islands in Galapagos and ancient connections among Neotropical dry forests

**DOI:** 10.1101/2021.05.11.443682

**Authors:** Ivan L. F. Magalhaes, Adalberto J. Santos, Martín J. Ramírez

## Abstract

Event-based biogeographic methods, such as dispersal-extinction-cladogenesis, have become increasingly popular for attempting to reconstruct the biogeographic history of organisms. Such methods employ distributional data of sampled species and a dated phylogenetic tree to estimate ancestral distribution ranges. Because the input tree is often a single consensus tree, uncertainty in topology and age estimates are seldom taken into account, even when they may affect the outcome of biogeographic estimates. Even when such uncertainties are taken into account for estimates of ancestral ranges, they are usually ignored when researchers compare competing biogeographic hypotheses. We explore the effect of incorporating this uncertainty in a biogeographic analysis of the 21 species of sand spiders (Sicariidae: *Sicarius*) from Neotropical xeric biomes, based on a total-evidence phylogeny including a complete sampling of the genus. By using a custom R script made available here, we account for uncertainty in ages and topology by estimating ancestral ranges over a sample of trees from the posterior distribution of a Bayesian analysis, and for uncertainty in biogeographic estimates by using stochastic maps. This approach allows for counting biogeographic events such as dispersal among areas, counting lineages through time per area, and testing biogeographic hypotheses, while not overestimating the confidence in a single topology. Including uncertainty in ages indicates that *Sicarius* dispersed to the Galapagos Islands when the archipelago was formed by paleo-islands that are now drowned; model comparison strongly favors a scenario where dispersal took place before the current islands emerged. We also investigated past connections among currently disjunct Neotropical dry forests; failing to account for topological uncertainty underestimates possible connections among the Caatinga and Andean dry forests in favor of connections among Caatinga and Caribbean+Mesoamerican dry forests. Additionally, we find that biogeographic models including a founder-event speciation parameter (“+J”) are more prone to suffer from the overconfidence effects of estimating ancestral ranges using a single topology. This effect is alleviated by incorporating topological and age uncertainty while estimating stochastic maps, increasing the similarity in the inference of biogeographic events between models with or without a founder-event speciation parameter. We argue that incorporating phylogenetic uncertainty in biogeographic hypothesis-testing is valuable and should be a commonplace approach in the presence of rogue taxa or wide confidence intervals in age estimates, and especially when using models including founder-event speciation.

## Introduction

The reconstruction of the biogeographic history of organisms is one of the main aims of systematic biology. Based on a known phylogeny, researchers may attempt to glimpse into the ancestral area where a particular clade originated (Huelsenbeck & Imennov 2002), infer the number and direction of dispersal events (Dupin et al. 2017), or estimate the number of vicariant events along the evolutionary history of the group (Ronquist 1997). More often than not, the geographic distribution of a group is known only from its extant species, occasionally accompanied by a few fossil forms. This represents but a fraction of the diversity of a group throughout its history, and we cannot directly assess the distribution range of unknown extinct species. Thus, biogeographers interested in such questions must resort to methods that attempt to estimate past distribution ranges from the data available from known species.

Initially, such estimates of ancestral geographical ranges relied on algorithms such as Fitch or Camin-Sokal parsimony optimization (Bremer 1992; Huelsenbeck & Imennov 2002). This approach treats distribution ranges as discrete characters and optimizes them as such along the phylogenies, but has some drawbacks. For instance, only tips of the phylogeny may be ‘polymorphic’ and occur in more than one area simultaneously, and important biogeographic processes such as vicariance are not modeled at all. This changed with the advent of event-based biogeography methods, pioneered by dispersal-vicariance analysis (DIVA; Ronquist 1997). Such methods attempt to explain the current distribution of organisms by modeling biogeographic events such as dispersal (colonization of a new area), range contractions (extinction of a species in a particular area) and vicariance (allopatric speciation leading to each descendant inheriting only part of the range of the ancestor species). DIVA brought a fundamental advance with respect to previous methods: changes in a lineage’s geographic distribution may happen as anagenetic events (dispersal or extinction along branches) or cladogenetic events (vicariance at nodes). Furthermore, the states in such estimates are geographic ranges, which may consist of one or more areas; thus, both tips and ancestors may be ‘polymorphic’. By using a parsimony framework, DIVA assigns costs to dispersal and extinction events, and treats vicariance as the null expectation for explaining biogeographic history. A similar rationale was used later to elaborate a likelihood-based approach to ancestral range estimation named dispersal-extinction-cladogenesis analysis (DEC; Ree et al. 2005; Ree & Smith 2008). DIVA’s parsimony-based optimization is agnostic regarding branch lengths. On the other hand, DEC models anagenetic events as a time-continuous function that takes branch lengths into account; thus, longer branches are more likely to contain anagenetic events such as dispersal or extinction. More importantly, DEC can accommodate dated trees, and thus prior information on geological history can be provided for explicit testing of biogeographic hypothesis (Ree et al. 2005). Modifications of DIVA and DEC have been implemented and further elaborated in software packages for biogeography, such as RASP (Yu et al. 2015) and BioGeoBEARS (Matzke 2013a, 2013b). The latter has become particularly popular due to its flexible implementation of biogeographic models allowing for different types of cladogenetic events, such as vicariance, subset sympatry, and founder-event speciation (see Matzke 2013b, 2014); in addition, all models are implemented in a likelihood framework, allowing direct model comparison.

A unifying feature of all methods discussed above is that they rely on the knowledge of the geographic distribution of each of the taxa, and of the phylogenetic tree describing their interrelationships. As in any comparative method, estimates of ancestral ranges cannot be more reliable than the data underlying it. The knowledge of the geographic distribution of tips may be affected by the Linnean and Wallacean shortfalls (Hortal et al. 2015), i.e., gaps in the data about existing species and their distributions. However, it is arguable that estimates of ancestral ranges are usually carried out by systematists that specialize in a particular taxon, who strive to include all or most known species and distribution records in their sampling, so as to mitigate any negative effects of such shortfalls. On the other hand, the knowledge of the underlying phylogenetic tree can be potentially more problematic. In recent years, the accumulation of genomic-scale studies has shown that the phylogenetic relationships of some clades are elusive even with massive quantities of data due to e.g., incomplete lineage sorting and/or very short internodes (e.g., Suh 2016; Ballesteros & Sharma 2019), and some portions of such trees are shrouded in topological uncertainty. There is uncertainty not only regarding the topology, but also in the estimates of divergence times. This uncertainty in age estimates is also resistant to the accumulation of genetic markers (see Ho & Duchêne 2014). Rates of molecular evolution rarely conform to a strict molecular clock, and branch lengths estimated from molecular sequences are a product of substitution rate and time, such that each of these two parameters are individually non-identifiable (Drummond et al. 2006). In addition, estimation of dated trees builds upon several assumptions, some of which may be too daring for empirical datasets (Bromhan 2019), and require fossil or other external calibrations, whose selection and justification is not free of difficulties (Parham et al. 2012, Warnock et al. 2015). In this scenario, analyses that depend on phylogenetic trees might benefit from accounting for any uncertainties regarding their topologies and/or ages (Huelsenbeck et al. 2000). In this paper, we refer to “phylogenetic uncertainty” as any uncertainty regarding the topology and/or node ages.

Efforts have been made to incorporate such phylogenetic uncertainty into ancestral range estimation. Nylander et al. (2008) used multiple trees from the posterior distribution of a Bayesian analysis to estimate ancestral ranges, and thus infer the biogeographic history of a clade of birds whose phylogeny had proven difficult to solve; they called this approach Bayes-DIVA. Similar pipelines have been incorporated into RASP (S-DIVA; Yu et al. 2010) and BioGeoBEARS (using the *run_bears_optim_on_multiple_trees* function). These functions summarize estimates of ancestral ranges of several trees on a target tree, such as a majority-rule consensus (as output by e.g., MrBayes) or a maximum clade credibility tree (MCC tree; as output by e.g., BEAST) and are routinely employed by researchers interested in incorporating phylogenetic uncertainty (e.g., Baker et al. 2020, Santaquiteria et al. in press). However, the results are summarized in a single tree, and the underlying variability in the estimates is difficult to grasp. Furthermore, the individual underlying trees usually are not taken into account when comparing competing biogeographic hypotheses. We argue that it is important to understand the effect of analyzing individual trees during hypothesis-testing, especially if confidence intervals of age estimates cross boundaries of time slices in time-stratified analyses.

In addition to phylogenetic uncertainty, there is the uncertainty associated with stochastic time-continuous models such as DEC. Because of this, estimates for ancestral nodes might include several possible states, each with its own probability. This hampers counting biogeographic events or estimating their ages. Dupin et al. (2017) solved this by introducing biogeographic stochastic mapping (BSM), a method to count biogeographic events while accounting for uncertainty in ancestral range estimation. Several replicates are run, and in each one the states at each node are resolved by taking into account the probabilities of each state. This allows, for instance, counting the number of dispersal events among areas, or estimating the time when such transitions took place. This approach is elegant, but it is usually employed on estimates based on a single tree, and thus incorporates uncertainty on ancestral range estimates conditional on a single topology. If there is topological uncertainty leading to different ancestral range estimates, this approach will underestimate the uncertainty in the biogeographic reconstruction (Fig. 1). Furthermore, using a single tree as input ignores the confidence intervals in age estimates, which are usually large. We argue that it would be productive to run stochastic maps over a sample of trees to incorporate both phylogenetic and stochastic uncertainty simultaneously.

**Figure 1.**
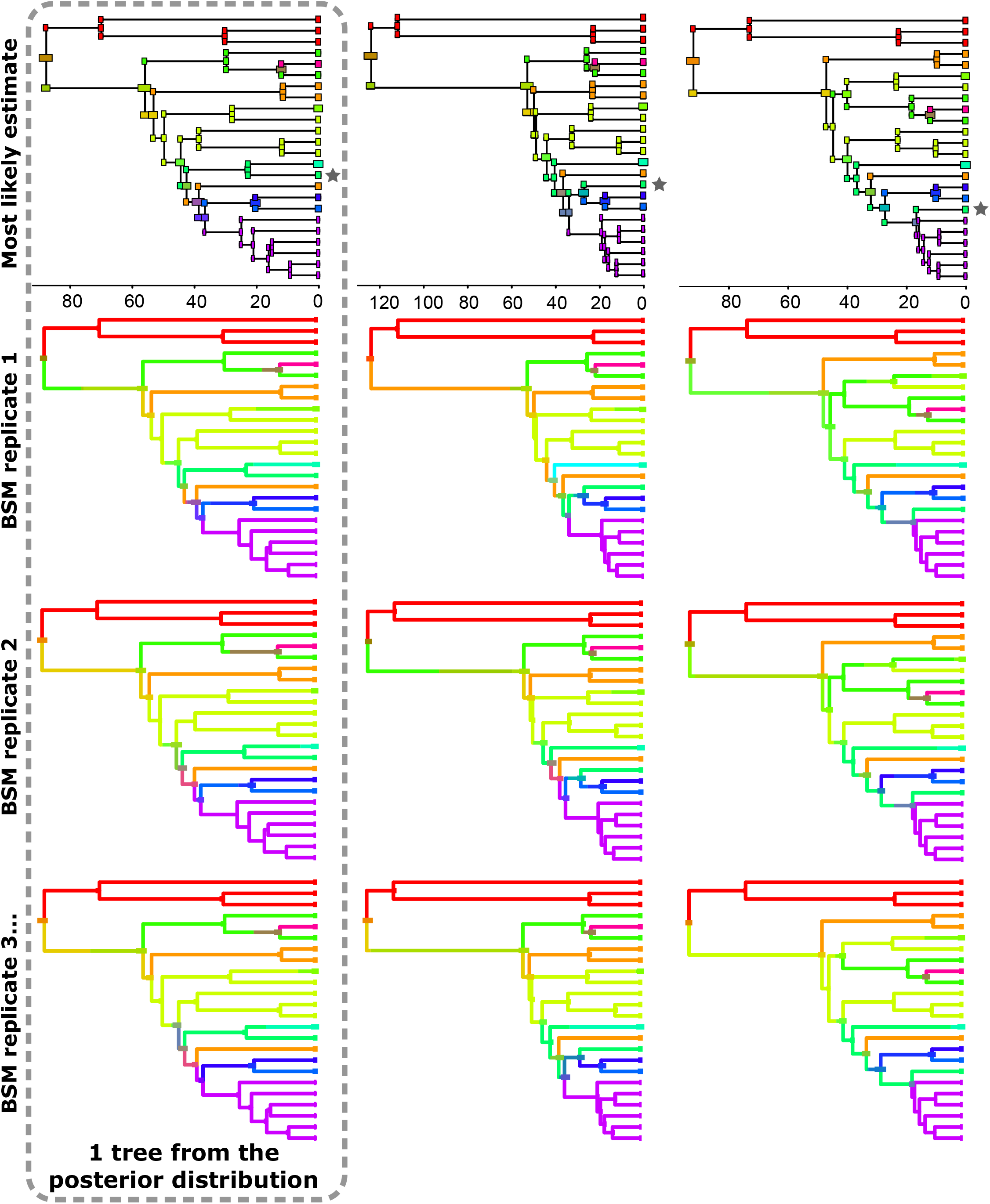
Biogeographic stochastic maps are insufficient for fully accounting for uncertainty in ancestral range estimates. Each column represents a single tree from the posterior distribution of the Bayesian analysis of our dataset, with the most likely estimated states in the top row, and different stochastic maps in the bottom rows. While estimates in each stochastic map are different, the most remarkable differences are found among trees different topologies. Note the tip marked with a grey star, which is a rogue taxon (*Sicarius andinus*). As it shifts its position across the different trees of the stationary phase of the chain, it leads to substantially different ancestral range estimates in each tree.

To illustrate our argument, we explore the effect of uncertainty in the biogeographic history of Neotropical sand spiders (*Sicarius*). These spiders represent an ideal system for this test because they are moderately diverse (21 species) and all species have been included in a dated total-evidence phylogeny (Magalhaes et al. 2019). The genus has a disjunct distribution, and each species is restricted to one or two arid areas surrounded by mesic habitats; phylogeographic and phylogenetic patterns suggest they are very poor dispersers (Binford et al. 2008; Magalhaes et al. 2014, 2017). Thus, their areas of distribution are clearly delimited and could be interpreted as “islands” of dry biomes inserted in a matrix of unsuitable humid habitats. Because of their distribution and moderate diversity, most of the biogeographic transitions should be straightforward to interpret, thus allowing us to easily measure the effects of phylogenetic uncertainty in biogeographic estimates.

Specifically, we focus on two particularly pressing questions. The first is the timing of colonization of the Galapagos archipelago by *Sicarius*. The Galapagos are currently inhabited by a single sand spider, *Sicarius utriformis* (Butler), which is sister to *S. peruensis* (Keyserling) from the Peruvian coastal deserts (Magalhaes et al. 2017, 2019). Although the oldest emerged islands are ∼3.5 million years (Myr) old (White 1993), geological evidence suggests that the archipelago existed for at least 14.5 Myr, when it was formed by paleo-islands that are now drowned (Christie et al. 1992; Werner et al 1999). The age of divergence between *S. utriformis* and *S. peruensis* has a 95% confidence interval between 1.2 and 22.2 Myr (median 9.7; Magalhaes et al. 2019), and thus it is possible that this pair of species split before the current islands were formed, but during the time the archipelago was formed by paleo-islands. This makes this system ideal to test the effect of the uncertainty of age estimates in biogeographic inference.

The second question is the estimation of the ancestral distribution range of a clade endemic to the Brazilian Caatinga. This area is one of the largest and most diverse tropical dry forests in the world (DRYFLOR 2016), and six *Sicarius* species inhabit the region (Magalhaes et al. 2017). These six species form a well-supported monophyletic group (Magalhaes et al. 2019), suggesting a single colonization of this area. In this case, from which area did the ancestor of this clade come? The American tropical dry forests and other xeric biomes such as deserts and scrublands currently have a disjunct distribution, but the similarity in their biota suggests they have been connected in the past (see DRYFLOR 2016). Basing on plant distributions, some argued that the Caatinga could have been connected to dry forests in Bolivia and Argentina, or to dry forests in the Caribbean coast of northern South America (Prado & Gibbs 1993; Pennington et al. 2000); these connections would have taken place by expansion of dry forests over areas that now are covered by mesic biomes. Connections between the Caatinga and the Caribbean dry forests imply a northern route passing through present-day Amazon (a rainforest); alternatively, the Caatinga could have been connected with southern formations, such as the Monte or the Chiquitano dry forests, passing through present-day Cerrado (a savannah). We thus aim at identifying the most likely route for the occupation of the Caatinga by sand spiders. However, this is hampered by the fact that one species, *Sicarius andinus* Magalhaes et al. from the Peruvian Andes, is a rogue taxon in the phylogeny and does not have a well-resolved phylogenetic position (Magalhaes et al. 2019). Different positions of this species may yield different biogeographic reconstructions for the Caatinga clade, and thus we must take this into account.

In this paper, we test the effect of taking phylogenetic uncertainty into account in the biogeographic inferences of the two pragmatic scenarios mentioned above. We combine processing of several trees of the stationary phase of the Markov chains of a Bayesian analysis with biogeographic stochastic maps for each tree, so that both phylogenetic and biogeographic uncertainties are considered simultaneously.

Specifically, we test (1) whether data from sand spiders support dispersal to Galapagos in the last 3.5 Myr (age of oldest emerged island), in the last 14.5 Myr (age of oldest known drowned paleo-island) or an unconstrained model (representing the possibility of older, yet undetected paleo-islands), and (2) whether the Caatinga was connected to northern (Caribbean, Mesoamerican or Andean dry forests) or southern (Chiquitano dry forests or Monte) biomes, as well as the age of such connections. We anticipate that analyzing a sample of trees, instead of a single target tree, provides invaluable insights for testing competing biogeographic hypotheses. Finally, we provide scripts for R (R Core Team 2020) to replicate the analyses described below, in the hope they will be useful for further studies.

## Material and Methods

### Summaries of biogeographic inferences in face of uncertainty

We prepared an R script to (1) sample *n* trees randomly from BEAST output files and prune them to the taxa of interest, (2) estimate ancestral ranges using BioGeoBEARS for each of these trees, (3) run biogeographic stochastic maps for each of these estimates, and finally (4) parse the results and summarize them. The summaries include: (1) parameter estimates, log-likelihoods and AICc for ancestral range estimates of each tree, (2) tables containing the transitions between geographic ranges for each stochastic map of each of the *n* sampled trees, along with the age of such transitions; (3) a graphic of lineages through time per area averaged over all trees and stochastic maps; (4) tables with all possible geographic ranges in both rows and columns, and average counts of how many times a transition between a particular pair of ranges took place; such tables are broken down by each type of transition (dispersal, extinction, vicariance, etc.) and summarized in a table containing all types of transitions; and (5) a summary of the most common biogeographic transitions (as the mean number of transitions per BSM replicate). The script (Online Supplementary File S1) uses functions from the packages phytools (Revell 2012), ape (Paradis & Schliep 2019), sjmisc (Lüdecke 2018) and BioGeoBEARS (Matzke 2013a) and has been written as to be easily adapted to most datasets. The necessary input files are the same as those used in BioGeoBEARS, except that users may provide multiple trees instead of a single target tree.

### Model selection

BioGeoBEARS implements three different biogeographic models that differ mainly in the events that may take place during cladogenesis when the ancestor has a widespread range (i.e., its range consists of two or more areas; see Matzke 2013b for a summary). DEC has an identical implementation to the model described by Ree & Smith (2008) and allows narrow vicariance (one descendant inherits exactly a single area, the other inherits the rest of the range) or subset sympatry (one descendant inherits exactly a single area, while the other inherits the whole distribution range of the ancestor). DIVA-like (DVL) allows the same cladogenetic events as the original implementation of DIVA (Ronquist 1997): narrow vicariance (as in DEC) and wide vicariance (each descendant may inherit two or more areas from the ancestor). BAYAREA-like (BAL) does not allow changes in distribution ranges during cladogenesis, and each descendant inherits exactly the same range as the ancestor. Each of these three models can be modified by the addition of a “jump-dispersal” free parameter (“+J”) that allows for founder-event speciation during cladogenesis, i.e., one of the descendants occupies a single area that is not part of the range of the ancestor, while the other descendant inherits the same range as the ancestor (Matzke 2014). Thus, models including founder-event speciation are unique in that they allow dispersal to take place during cladogenetic events.

Biogeographic history may be estimated under each of these six models and the fit of the data to each of them can be compared by using the Akaike information criterion (AIC; Akaike 1973) and Akaike weights (AICw; Wagenmakers & Farrell 2004). This procedure has been used to guide the selection of models that best explain the data and for testing biogeographic hypotheses (Matzke 2013b, 2014). We here estimate the fit of the data to six different models (DEC, DVL, BAL, DEC+J, DVL+J, BAL+J) for initial exploration of the behavior of the estimates. The script used for model selection can be found as Online Supplementary File S5.

We wanted to investigate whether the choice of a particular biogeographic model interacts in any way with the decision to run analyses over a sample of posterior trees. Thus, we ran the aforementioned script under two models with (DIVA-like+J and BAYAREA-like+J) and without (DIVA-like and DEC) founder-event speciation. The choice of these four models was done to include those that are a better fit to the data (DIVA-like and the +J variant), a model that is frequently used in empirical studies (DEC) and a model excluding the possibility of vicariance, and thus more similar to simple parsimony reconstructions (BAYAREA-like+J). Each of these four models was used in both unconstrained and time-stratified analysis (see below), and using either the MCC tree or 100 posterior trees as source. In this latter case, to make these analyses directly comparable, the same sample of 100 trees was used for all runs. The combination of four biogeographic models, three time-stratification scenarios and two sources of trees resulted in 24 different runs (see Online Supplementary Figure S10).

### Phylogeny and distribution data

To reconstruct the biogeographic history of *Sicarius*, we used a recently published phylogeny estimated using morphology and DNA sequences and dated using a combination of fossil calibrations and substitution rates for the histone H3, subunit A gene (Magalhaes et al. 2019). To understand the effect of incorporating uncertainty into biogeographic inference, analyses were run both on the maximum clade credibility (MCC) tree, and on a sample of 100 trees randomly drawn from the posterior distribution of the analysis, after removing the first 10% samples as burn-in (see above). *Hexophthalma* is the African sister group of *Sicarius* and was included in the analyses; the remaining terminals (*Loxosceles* and non-sicariid outgroups) were pruned from the trees. Species geographic ranges and trees can be found as Online Supplementary Files S2–S4.

The distribution of each species has been fully mapped in recent taxonomic publications on the genus including *ca*. 1800 adult specimens from natural history collections and recent field expeditions (Cala-Riquelme et al. 2017; Magalhaes et al. 2017). We classified species distribution in ten areas: southern Africa deserts and xeric scrublands (F), Argentinean M<u>o</u>nte (O), Atacama Desert and neighboring Chilean xeric scrublands (T), <u>S</u>echura desert in the Peruvian coast (S), An<u>d</u>ean dry forests (D), <u>C</u>hiquitano dry forests in Bolivia (C), <u>M</u>esoamerican dry forests (M), dry forests in the Cari<u>b</u>bean coast of Colombia (B), Caatinga dry forest in Brazil (I), and the <u>G</u>alapagos Islands (G) (Fig. 2). Most of these areas are clearly delimited by geographic barriers such as oceans or mountain ranges, or correspond to well-recognized ecoregions or phytogeographic units (DRYFLOR 2016; Echeverría et al. 2018).

**Figure 2.**
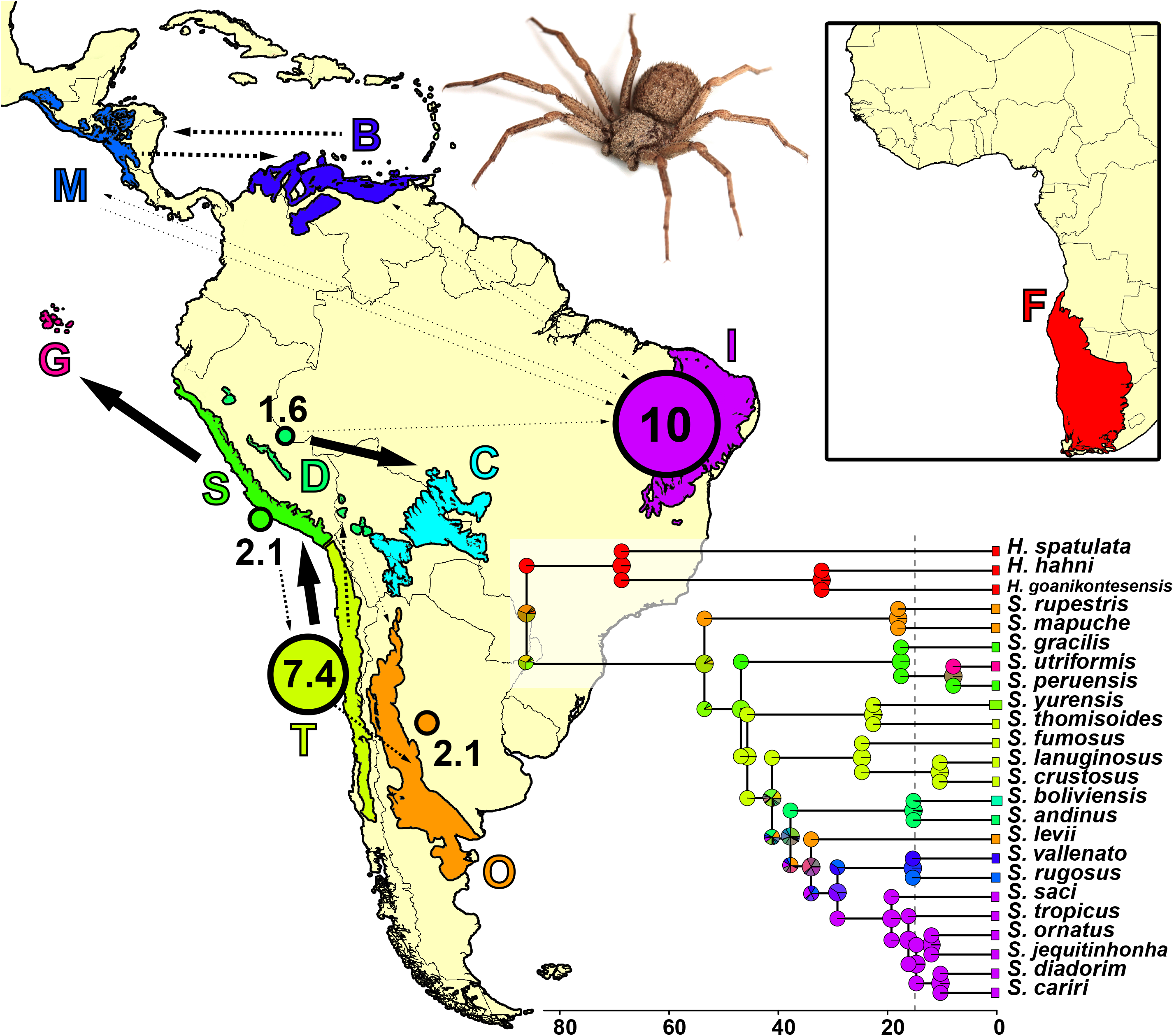
Map depicting the areas inhabited by *Sicarius* and *Hexophthalma*, and the ancestral range estimates under DIVA-like using the maximum clade credibility tree. Solid arrows among areas represent one dispersal event between areas that is robust to topological and biogeographical uncertainty. Dashed lines represent inferred dispersal events that are sensitive to uncertainty; arrow widths are proportional to the frequency at which such dispersals are inferred. Numbered circles indicate the inferred number of within-area speciation in each of the areas. Area abbreviations: B = dry forests in the Cari<u>b</u>bean coast of Colombia; F = southern Africa deserts and xeric scrublands; C = <u>C</u>hiquitano dry forests in Bolivia; D =An<u>d</u>ean dry forests; G = <u>G</u>alapagos Islands; I = Caatinga dry forest in Brazil; M = <u>M</u>esoamerican dry forests; O = Argentinean M<u>o</u>nte; S = <u>S</u>echura desert in the Peruvian coast; T = Atacama Desert and neighboring Chilean xeric scrublands. Africa not to scale.

### BioGeoBEARS parameters and time stratification

Sand spiders are poor dispersers (Magalhaes et al. 2019) and most species are restricted to a single area, with only two species occurring in two areas (Magalhaes et al. 2017). For this reason, and to speed up calculations, we restricted the maximum range size to include three areas. Likelihood calculations were carried out with *optimx* (Nash 2014). We compared ancestral ranges estimates under three scenarios: (1) unconstrained, allowing dispersal to the islands in any time, and (2 & 3) two different time-stratified scenarios, each with two time slices, where the Galapagos Islands were only available for occupation in the more recent slice. The boundary between the two slices was set to (2) 15 Myr, as geological evidence points to drowned islands that are at least 14.5 Myr old (Werner et al. 1999), or (3) 3.5 Myr, representing the approximate age of the oldest emerged island (White et al. 1993). Files for implementing the time-stratified model are available as Online Supplementary Files S6–8.

## Results

### Model selection and estimates of ancestral ranges

Regarding runs on MCC trees, overall DEC and DIVA-like resulted in similar ancestral range estimates among them, as did all models including a +J parameter. Parameter estimates and fit of data under different models are summarized in Online Supplementary Table 1. AIC values indicate that DIVA-like (log-likelihood: -51.29, AIC: 107.16) is the favored model among those not including a founder-speciation free parameter, while DIVA-like + J (log-likelihood: -44.24, AIC: 95.69) is favored among models including this parameter. Because it has been demonstrated that models including founder-event speciation are prone to over-fitting (Ree & Sanmartín 2018; but see Matzke 2021), we here show results of the ancestral range estimates for the MCC tree under DIVA-like (Fig. 2); results under DIVA-like+J can be seen in Online Supplementary Figure 11.

**Table 1.** Geographic range of origin of clades dispersing into the Caatinga and their relative frequencies (in %) in different biogeographic stochastic maps. Taking topological uncertainty into account (by running stochastic maps in 100 posterior trees) increases the percentage of inferences of dispersal coming from Andean or Chiquitano dry forests (values marked in bold) while decreasing the probability of dispersal coming from Mesoamerican or Caribbean dry forests, or the Atacama desert (values marked in italics).

| Source trees<br>(each with<br>100 BSM) | Biogeographic<br>model | Caribbean (B) | Chiquitano (C) | Andes (D) | Mesoamerica (M) | Monte (O) | Atacama (T) | Caribbean+Andes | Andes+Mesoamerica | Caribbean+Mesoamerica | Caribbean+Monte | Andes+Monte | Monte+Mesoamerica |
| --- | --- | --- | --- | --- | --- | --- | --- | --- | --- | --- | --- | --- | --- |
| MCCT | DVL+J | 25 | 0 | 13 | 30 | 15 | 15 | 0 | 0 | 2 | 0 | 0 | 0 |
| MCCT | DVL | 22 | 0 | 9 | 19 | 19 | 19 | 0 | 1 | 2 | 2 | 4 | 3 |
| 100 trees | DVL+J | 22 | **2** | **24** | 22 | **16** | *11* | 0 | 0 | 2 | 0 | 0 | 0 |
| 100 trees | DVL | *16.2* | **1.2** | **26.6** | *15.3* | *12.5* | *15.5* | **1.3** | 1.3 | 2.7 | 1.9 | *1.7* | 2 |

We briefly investigated the effect of taking topological uncertainty into account during model selection by comparing the AICc values of the estimates of each of the 100 trees for the models DIVA-like, DIVA-like+J and DEC. A histogram of such values displays some overlap among values of the different models (Online Supplementary Figure 12). However, for each individual tree the relationship of the preferred model is maintained (i.e., DIVA-like+J is preferred to DIVA-like, which is preferred to DEC), and is identical to the order found by running the analysis using the MCC tree. Thus, taking uncertainty into account has not affected the results of model selection.

### Lineages through time by area

We estimated the number of lineages occupying each individual area through time. For this, the tree has been divided in 1 Myr slices, and we counted the number of species in each area in each slice; widespread species are counted once in each area of their distribution range. The number of lineages in each area increases with time (Fig. 3) as a combination of within-area speciation and new dispersals into the area. When using a single tree, changes in diversity associated with cladogenetic events (e.g. within-area speciation or founder-event speciation) appear as abrupt increases in the plot (Fig. 3a, b). This effect is stronger in the model including founder-event speciation (Fig. 3a), but disappears when topological and age uncertainty is taken into account (Fig. 3c, d).

**Figure 3.**
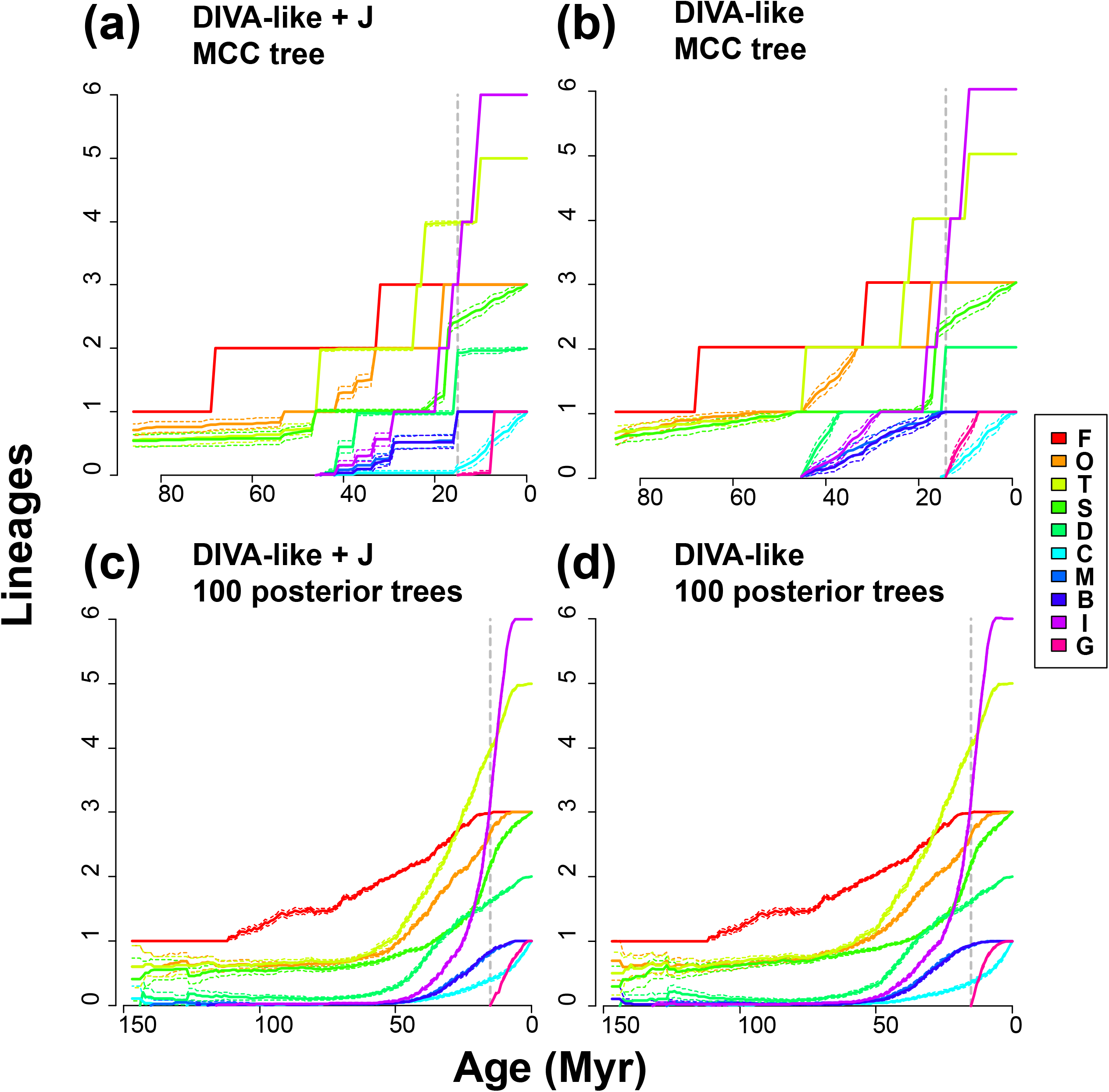
Number of lineages occupying each area through time. Solid lines are the average of 100 biogeographic stochastic maps, and dashed lines are the 95% confidence interval. See Fig. 2 for area abbreviations. The dashed vertical line indicates the boundary between time slices in the time-stratified analysis. Runs on a single maximum clade credibility tree (a, b) show abrupt changes in the number of lineages that are related to cladogenetic events, whose age does not vary because there is no uncertainty associated to the tree. This effect is more pronounced in the model with founder-event speciation (a) because it relies more on cladogenetic events for estimating ancestral ranges. Using several trees (each with 100 stochastic maps) to account for topological and age uncertainty removes this effect and smooths the curves (c, d), reducing differences between models with and without founder-event speciation.

The results (Fig. 3) indicate that the ancestor of *Hexophthalma* + *Sicarius* lived in a range composed of (1) southern African deserts and scrublands, and (2) either the Atacama desert, Sechura desert, or Argentinean Monte. These latter three have similar probabilities of being part of the ancestral range of sand spiders due to uncertainty in topology and biogeographic estimates. There has been a slow and steady increase in diversity in these temperate desert areas for the last ∼100–80 Myr. On the other hand, lineages only started occupying tropical dry forests later, around 50 Myr. The Caatinga has become the most species-rich region relatively rapidly due to within-area speciation, mainly during the Miocene.

### Comparing time-stratified *vs*. unconstrained models in the face of age uncertainty

We compared the fit of the data to unconstrained models (occupation of Galapagos possible at any time) to time-stratified models (occupation of Galapagos possible only in the last 15 Myr, or in the last 3.5 Myr). First, we investigated the inferred age of dispersal to the islands in the unconstrained analysis. In the analyses using a MCC tree as input, dispersal to the Galapagos has been inferred to occur more recently than 15 Myr in 72% (DIVA-like; mean 13.8 Myr) or 97% (DIVA-like+J; mean 8.8 Myr) of the stochastic maps (Fig. 4a, b). Using a MCC tree results in sharper, overconfident distributions of the inferred ages of dispersal to the islands. This is especially notable in the case of the model with founder-event speciation, where 88% of the replicates inferred that colonization of the islands is a cladogenetic jump-dispersal with age equal to that of the *S. utriformis–S*.*peruensis* node in the MCC tree (Fig. 4a). In analyses using 100 posterior trees as input, dispersal to the Galapagos has been inferred to occur more recently than 15 Myr in 57.8% (DIVA-like; mean 16 Myr) or 78.2% (DIVA-like+J; mean 12 Myr) of the replicates (Fig. 4c, d). Thus, when using 100 posterior trees, inferred ages of the dispersal to Galapagos are older in average. In addition, the distribution of inferred ages is flatter, as it takes into account the uncertainty in node ages; this reduces the difference between models with and without founder-event speciation.We then compared the fit of the data to unconstrained model *vs*. time-stratified model allowing dispersal only in the last 15 Myr. Using the MCC tree, the data fit better to a stratified model under DIVA-like, DIVA-like+J, and BAYAREA-like+J, while it fits better to an unconstrained model under DEC (stars in Fig. 5). In all cases, however, the support for the preferred model is very weak, as the ratio between AICc weights of the preferred model range only between 1.29 to 2.58, and thus we cannot decisively reject any of the two models in favor of the other. When comparing models over 100 posterior trees, we observed an interesting pattern. Again, in most trees the data fit slightly better a time-stratified scenario under DIVA-like, DIVA-like+J, and BAYAREA-like+J (in 58, 68 and 58 of the 100 trees, respectively), and an unconstrained scenario under DEC (in 91 of the trees), but relative supports in these cases are once again ambiguous (1.67–2.91). However, in the fewer cases of trees where the data fits better an unconstrained model, the median relative support is higher and shows decisive support for this model (5.00–151.74) (Fig. 5).

**Figure 4.**
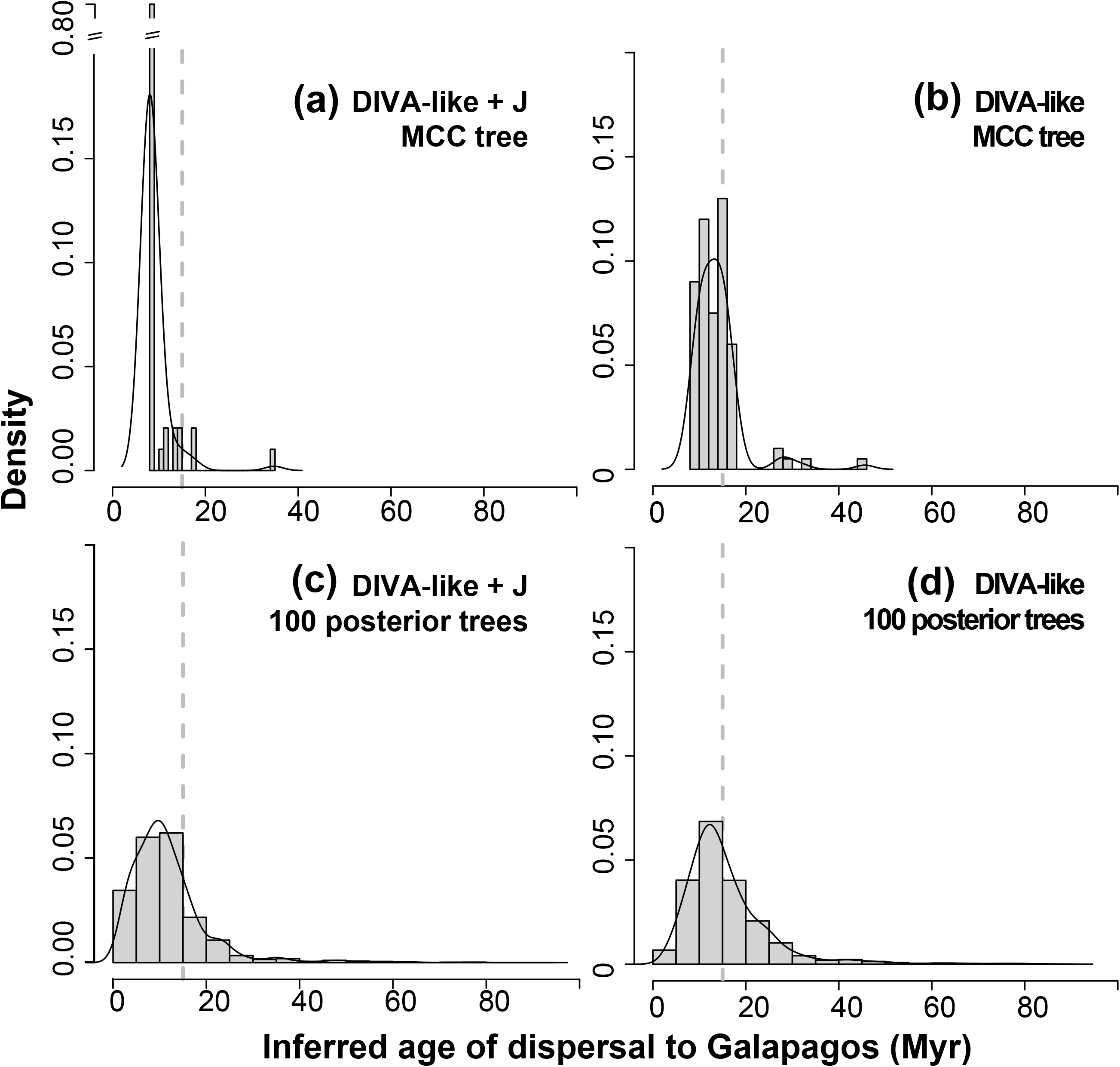
Histograms with the inferred age of dispersal of *Sicarius* to the Galapagos Islands. Analyses based on a maximum clade credibility tree (MCC tree) (a, b) produce sharper estimates that are closely tied to the age of split of *Sicarius utriformis* (from Galapagos) and its sister species. This effect is especially pronounced in the model including founder-event speciation (a), where 88% of the estimates have exactly the same age of that split. Accounting for uncertainty in topology and age estimates (c, d) reveals that the age of dispersal is much more uncertain. The dashed vertical line indicates the boundary between time slices in the time-stratified analysis and corresponds to the geological evidence of the oldest drowned paleoislands of the Galapagos archipelago.

**Figure 5.**
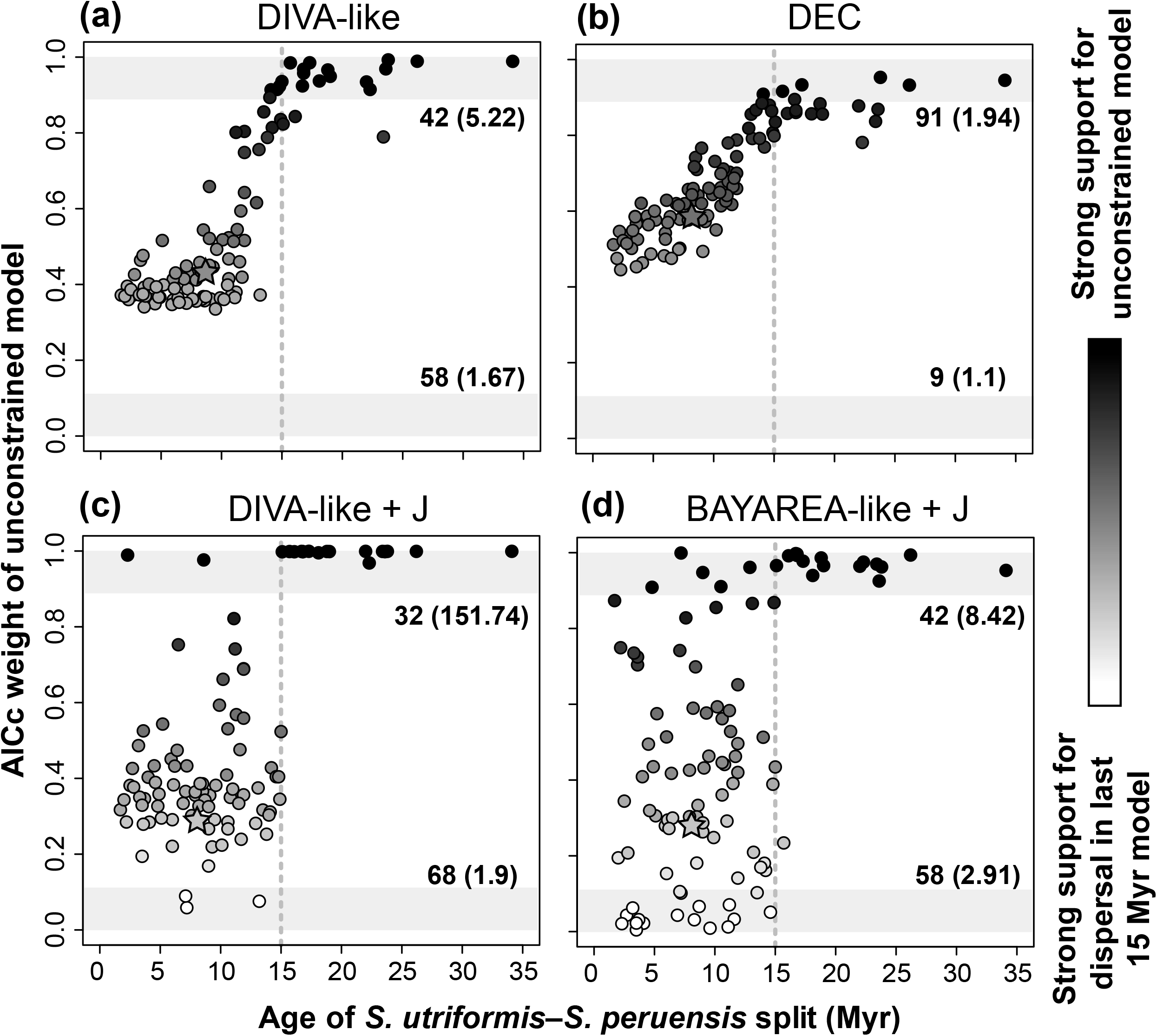
Relationship between AIC weight of the unconstrained biogeographic model allowing for dispersal of the Galapagos at any time (relative to the time-stratified model allowing dispersal only during the last 15 Myr) and the age of split of *Sicarius utriformis* (from Galapagos) and its sister species. Each dot represents an individual tree of a sample of 100 taken from the posterior distribution of a Bayesian analysis; the star represents the maximum clade credibility tree. Values of AIC weight close to 1 (darker shades) indicate strong support for the unconstrained model, while values close to 0 (lighter shades) indicate strong support for the time-stratified model. The light grey areas at the top and bottom of the graphs are the zones where the support for one of the alternative models is decisive. The dashed vertical line indicates the boundary between time slices in the time-stratified analysis. Numbers at the top and bottom of the graph are the number of trees supporting each model, and the median support relative to the alternative model. Trees in which the split between *S. utriformis* and its sister species are older than 15 Myr fit better to an unconstrained scenario allowing occupation of the Galapagos before that time.

We suspected that this pattern might be due to the age of split between *S. utriformis–S. peruensis*. The confidence interval of this age spans a wide range (1.2– 22.2 Myr) and actually crosses the boundary between the two time slices of the time-stratified model (15 Myr). To investigate this, we plotted the AICc weight of the unconstrained model against the age of the split (Fig. 5). The plot indicates that when the age of the split is younger than 15 Myr, there is weak support for the time-stratified model (DIVA-like and DIVA-like+J), to the unconstrained model (DEC), or the support to either of them is ambiguous (BAYAREA-like+J). On the other hand, the data fits better to the unconstrained model in all the 17 trees where the *S. utriformis–S. peruensis* split is older than the boundary of the time slice (15 Myr). While this is true for the four biogeographic models employed, the effect is much stronger in models including a founder-event speciation parameter, which tend to favor the unconstrained model more strongly: the relative support for the unconstrained model is higher in DIVA-like+J (776.2 ± 369) and BAYAREA-like+J (85.7 ± 110.86) than in DIVA-like (37.81 ± 37.97) and DEC (8.84 ± 4.73). Thus, in at least some of the trees from the posterior distribution, there is strong support for an unconstrained colonization of the Galapagos taking place before 15 Myr.

Finally, we compared the fit of the data to a model allowing dispersal to the Galapagos in the last 15 Myr (age of the oldest recorded paleo-islands) to a model allowing dispersal only in the last 3.5 Myr (age of the oldest emerged island). Under all biogeographic models, the model allowing dispersal in the last 15 Myr is strongly favored (Fig. 6). In the few trees where this split is younger than 3.5 Myr, the 15-Myr model is still strongly favored by DIVA-like and DEC, while support to either scenario is ambiguous in DIVA-like+J and BAYAREA-like+J. Thus, our data indicates that dispersal of sand spiders to the Galapagos took place before the appearance of the oldest emerged island.

**Figure 6.**
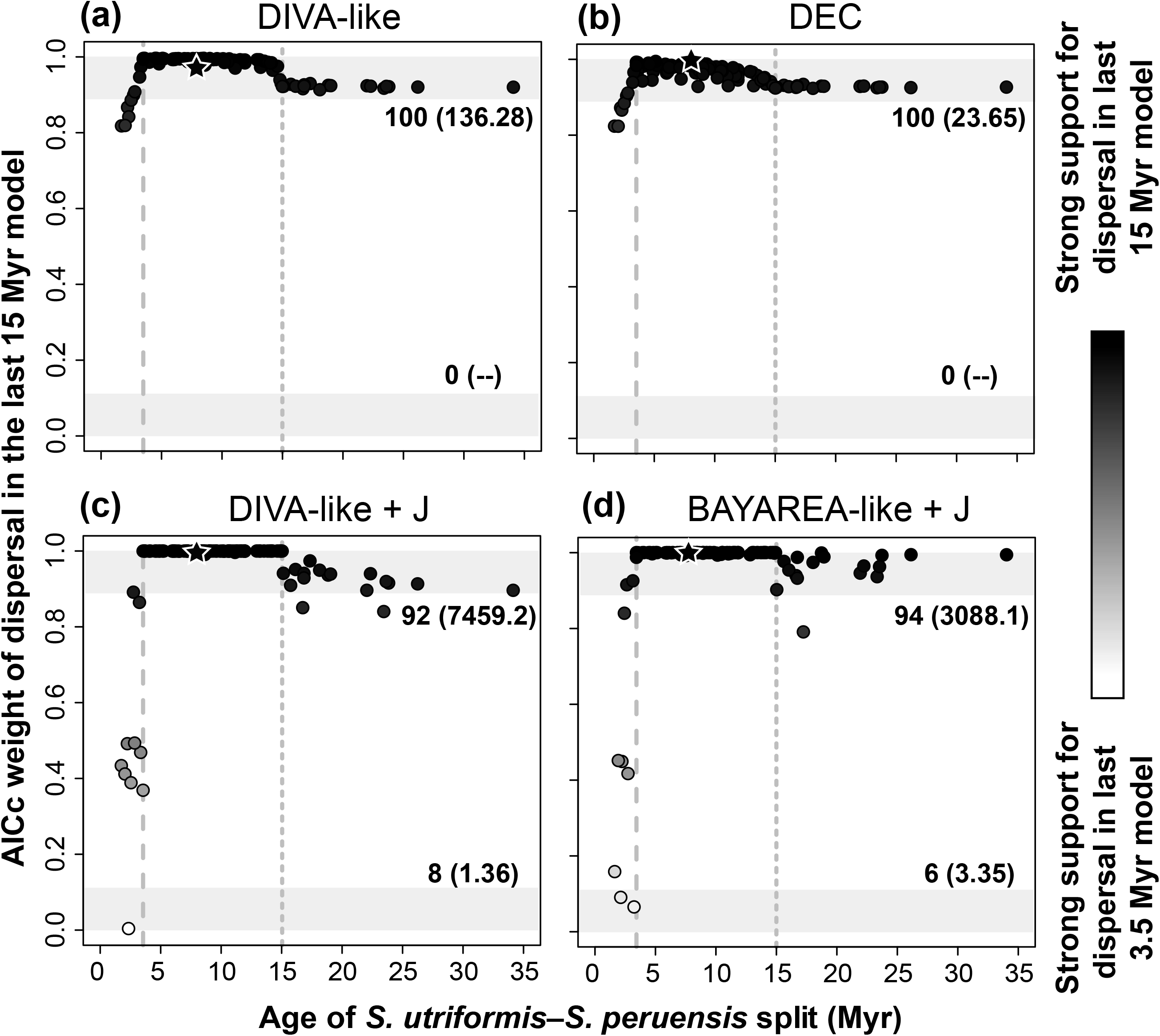
Relationship between AIC weight of the time-stratified biogeographic model allowing dispersal only during the last 15 Myr (relative to the time-stratified model allowing dispersal only during the last 3.5 Myr) and the age of split of *Sicarius utriformis* (from Galapagos) and its sister species. Each dot represents an individual tree of a sample of 100 taken from the posterior distribution of a Bayesian analysis; the star represents the maximum clade credibility tree. Values of AIC weight close to 1 (darker shades) indicate strong support for the 15 Myr model, while values close to 0 (lighter shades) indicate strong support for the 3.5 Myr model The light grey areas at the top and bottom of the graphs are the zones where the support for one of the alternative models is decisive. The dashed vertical lines indicate the boundary between time slices in each of the time-stratified models. Numbers at the top and bottom of the graph are the number of trees supporting each model, and the median support relative to the alternative model. With few exceptions, most trees fit decisively better to a model allowing for dispersal in the last 15 Myr, regardless of the age of the split.

### Ancestral range estimates of particular nodes in the face of topological uncertainty

We estimated the most likely state in the root node (corresponding to the split between African *Hexophthalma* and Neotropical *Sicarius*). For simplicity, we only report the results under DIVA-like, which are similar to those of other models. Analyses using the MCC tree yield estimates for an ancestral range most likely including southern African scrublands and one of the temperate American deserts (Sechura, Atacama or Monte): most likely ranges are FOT (30.7%), FOS (30.3%), FTS (24.6%), FO (4.8%), FT (3.8%) and FS (3.8%), summing to a total of 98.3%. The most likely ranges across the 100 trees are FOT (27.8%), FOS (24.4%), FTS (22.4%), FO (5.5%), FT (4.2%) and FS (3.7%), total 88.1%. These latter ranges have their likelihoods slightly diminished because of an increase in the likelihood of ranges including Andean dry forests, namely FOD (2.3%), FSD (2%), FTD (1.5%). At any rates, these changes are rather small, and, even in the face of topological uncertainty, we can be fairly confident that the ancestor of *Sicarius* + *Hexophthalma* lived in a range including deserts and xeric scrublands of southern Africa and southern or western South America.

We used stochastic maps to estimate the ancestral range that sourced species to the Brazilian Caatinga. In more than 99.9% of the maps, occupation of the Caatinga is the result of a single dispersal event, either jump-dispersal (DIVA-like + J) or anagenetic dispersal to a wide range including the Caatinga immediately followed by vicariance (DIVA-like). Using 100 stochastic maps resolved from the ancestral range estimates on the MCC tree, the most likely candidates for the area that originated this single dispersal are Mesoamerican dry forests, Caribbean dry forests, Argentinean Monte, Atacama desert or Andean dry forests (Table 1). When accounting for topological uncertainty using 100 stochastic maps for each of the 100 posterior trees, the results are similar but there is a substantial increase in the likelihood of Andean dry forests to be the source area, and this area actually becomes the most likely source (Table 1); using only the MCC tree underestimates this possibility. This increase in the likelihood is related to the presence of the rogue taxon *Sicarius andinus*, who inhabits Andean dry forests and is resolved as the sister taxon to the Caatinga clade in several trees of the posterior distribution.

We inferred the age of the dispersal event to the Caatinga and discovered a similar pattern to what we observed regarding the Galapagos. When using a MCC tree, the estimates are sharp and overconfident (Fig. 7a, b), especially in the analysis including founder-event speciation, which shows four peaks closely tied to the ages of the cladogenetic events immediately leading to the Caatinga clade. Using 100 posterior trees to estimate the age of the dispersal event incorporates the uncertainty in the node ages, and reduces the differences between DIVA-like and DIVA-like+J (Fig. 7c, d).

**Figure 7.**
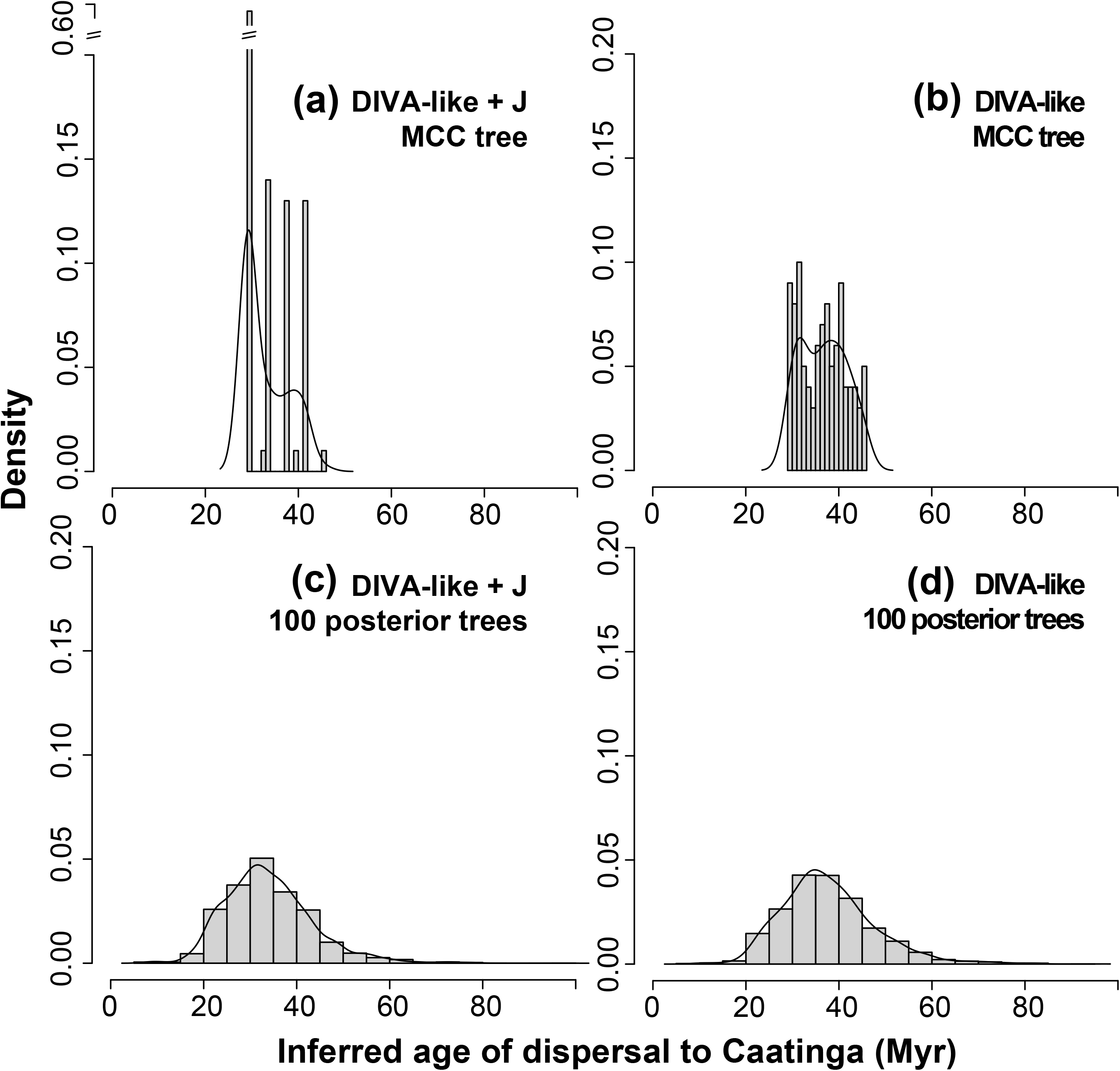
Histograms with the inferred age of dispersal of *Sicarius* to the Caatinga. Analyses based on a maximum clade credibility tree (MCC tree) (a, b) produce sharper estimates closely tied to the ages of cladogenetic events in this tree. This is especially notable in the model including founder-event speciation (a), which displays four peaks, each associated with the age of the four successive nodes in the tree that could have originated the founder-event dispersal to the Caatinga. Accounting for uncertainty in topology and age estimates (c, d) reveals that the age of dispersal is much more uncertain.

### Summary of biogeographic events in the face of uncertainty

We were able to count the most frequently inferred biogeographic events (dispersals, extinctions, vicariance and within-area speciation) by taking into account uncertainty in topology and ancestral range estimates. The results are summarized in Fig. 2 and Table 2. A substantial fraction of *Sicarius* diversity has been generated by within-area speciation, mainly in the Caatinga and the Atacama and, to a lesser extent, in the Monte scrubland, Sechura desert and Andean dry forests. Three of the dispersal events are robust to both types of uncertainty: one from the Sechura to Galapagos, one from the Atacama to Sechura, and one from the Andes to the Chiquitano dry forest. Other dispersal events are sensitive to uncertainty in topology and/or in ancestral range estimates, but generally involve geographically close areas, such as Mesoamerican and Caribbean dry forests, Atacama and Monte, and Atacama and Andes. The Caatinga has exchanged lineages with a single other area whose identity is uncertain, but candidates are Mesoamerican, Caribbean or Andean dry forests and, less likely, the Atacama desert and the Monte scrubland.

**Table 2.** List of the most common biogeographic events inferred under DIVA-like, averaged over 100 trees from the posterior distribution.

| Starting range | Ending range | Average events | Event type |
| --- | --- | --- | --- |
| I | I | 10.0948 | In-situ speciation |
| T | T | 7.5704 | In-situ speciation |
| F | F | 3.9997 | In-situ speciation |
| S | S | 2.5332 | In-situ speciation |
| O | O | 2.168 | In-situ speciation |
| D | D | 1.6566 | In-situ speciation |
| T | TS | 1.0111 | Dispersal |
| MB | M | 0.9475 | Vicariance |
| MB | B | 0.9475 | Vicariance |
| D | DC | 0.8726 | Dispersal |
| SG | S | 0.8004 | Vicariance |
| SG | G | 0.8004 | Vicariance |
| S | SG | 0.7997 | Dispersal |
| OT | O | 0.6377 | Vicariance |
| OT | T | 0.636 | Vicariance |
| TS | S | 0.5241 | Vicariance |
| TS | T | 0.5228 | Vicariance |

## Discussion

### Inference of biogeographic events in the face of uncertainty

In recent times, we have learned that tree topology may reach stability with more sequence data, but some clades remain elusive even in massive phylogenomic datasets (Suh 2016, Ballesteros & Sharma 2019). In addition, estimates of ages of divergence are based on limited data and many assumptions (e.g. the placement of fossil calibrations) and will always be uncertain; as Bromham (2019) neatly pointed out, “paleontological evidence and molecular dates paint history with a broad brush, not fine penwork”. In this scenario, it seems unwise to put too much faith in a single tree, which represents only one of many similarly probable topologies and age estimates. Our results corroborate this: when combining biogeographic stochastic maps with a sample of trees, there are valuable insights to be gained.

We illustrate this concept by studying particular biogeographic events. Biogeographic maps can be used to estimate both transitions among areas and ages of biogeographic events (Dupin et al. 2017). However, they are tied to the particular tree that is used as a base, which may bias the biogeographic inferences. When we estimated biogeographic transitions between the Caatinga and other dry Neotropical areas, analyses using only the MCC tree underestimated the possibility of dispersal coming from the Andes when compared to the analyses using 100 posterior trees (Table 1). Perhaps more importantly, the inferred ages of biogeographic events are severely biased when stochastic maps are run on a single tree (Figs 3a, b, 4a, b, 7a, b). This is because stochastic maps can only infer the ages of events along the branches and nodes of that particular tree. This bias is potentially problematic, because many biogeographic studies are aimed at linking phylogenetic and geological history; the correlation (or lack thereof) of node ages (or biogeographic events) with geological events is often used to reach biological conclusions (e.g. Renner 2016). If such uncertainty is not taken into account, researchers may achieve inaccurate results. We show that running stochastic maps on a single tree does not fully capture all the possible biogeographic histories of a clade, and thus strongly encourage researchers to use this method in combination with a sample of trees to take advantage of its full potential.

Additionally, it seems that using several trees reduces the differences between different biogeographic models (e.g. DIVA-like and its + J variant). We find that lineages through time by area and inferred ages of dispersal are more similar among models when phylogenetic uncertainty is taken into account (Figs 3c, d, 4c, d, 7c, d). Thus, at least in some cases, phylogenetic uncertainty might be more influential than uncertainty in the choice of a particular biogeographic model.

On a lighter note, our results suggest that phylogenetic uncertainty is not crucial for model selection. We show that the outcome of model selection (DEC, DIVA-like, BAYAREA-like, and their +J variants) is the same regardless if the comparison is done using the MCC tree or 100 posterior trees (Online Supplementary figure S12). Thus, it seems that model selection can be done using the MCC tree, and biogeographic stochastic maps can be run on a sample of posterior trees using only the preferred model.

### Cladogenetic events, founder-event speciation, and uncertainty

We find that bias in age estimates is stronger for cladogenetic events (relative to anagenetic events) when a single tree is used. Models including founder-event speciation disproportionately favor cladogenetic events (Ree & Sanmartín 2018), and thus are more prone to overconfidence, especially when inferring ages of biogeographic events. Such models have been introduced by Matzke (2013b, 2014) as a way to model the possibility of dispersal to a new area being followed by speciation over the course of very few generations—almost instantaneously in an evolutionary timescale. In practice, this parameter allows dispersal to happen simultaneously with cladogenetic events, i.e. at tree nodes. This is opposed to anagenetic dispersal, which happens along the branches. Stochastic maps resolve the age of an anagenetic dispersal at any point of such a branch, but founder-event dispersal is always tied to the age of a particular node. Because of this, ages of events inferred from models including founder-event speciation are more biased and closely tied to the ages of nodes when a single tree is used to run the stochastic maps (Figs 3a, 4a, 7a). This overconfidence is partly smoothed by the variability in ages that can be incorporated by using a sample of posterior trees (Figs 3c, d, 4c, d, 7c, d). Thus, we recommend incorporating phylogenetic uncertainty when running stochastic maps especially when using models including founder event-speciation. It is especially important to have this in mind since model selection often favors such models (Matzke 2014, Ree & Sanmartín 2018, Matzke 2021). It should also be noted, however, that even models favoring anagenetic dispersal also suffer from bias when using a single tree: they are bound to a given time interval, as opposed to the wider range of possible time intervals when several trees are used.

### Lineages through time by area

We here explore the concept of “lineages through time by area” plots. These were initially conceived by Ceccarelli et al. (2019), who were interested in tracking changes in diversity of two different areas through time. By dividing the tree into user-defined time slices, it is possible to count the number of lineages occupying each area in each time period. Diversity increases through time as a combination of in-situ speciation and new dispersals into the area. Additionally, diversity can also decrease if the model infers extinctions in a particular area (Supplementary Figure S13), or if the taxon sampling includes fossil tips (ILFM, unpublished data).

These plots may be used as rough approximations to detect areas which might be under special processes. For instance, our data on sand spiders indicate that the Caatinga rapidly accumulated diversity after the initial dispersal to this area (Fig. 3), which is consistent with observations that this is one of the most diverse dry forests in the Neotropical region (DRYFLOR 2016). Nevertheless, it should be noted that these plots present limitations. It is known that DEC and its variants vastly and consistently underestimate extinctions (Ree & Smith 2008). Known but unsampled species are effectively “extinct” for the purposes of the model and will not be considered, which may affect the results of the plot, particularly if the sampling is uneven across areas. This is clear when we take a look at the diversification of lineages from Africa: only four in-situ speciation events are inferred (Fig. 3, Table 2), even though the genus *Hexophthalma* currently includes 8 species (Lotz 2018). It is clear that diversification in this area is underestimated by our sampling, which only includes three species. Thus, we recommend that such plots are only considered when the sampling of the group is complete, or at least when the unsampled species are not biased to a particular area. Second, even if the sampling is complete, these plots will not account for truly extinct species. Researchers interested in comparing diversification rates among areas, while accounting for extinct lineages, should refer to methods specifically designed for modelling this (e.g. BAMM; Rabosky 2014).

### Sand spiders dispersed to Galapagos paleo-islands

The Galapagos Islands are volcanic in origin, and while the oldest emerged island is 3–4 million years old (White et al. 1993), there is evidence of drowned paleo-islands to the east (Christie et al. 1992) and north (Werner et al. 1999) of the current archipelago. The particular biota of the islands has affinities with those of South America, the Greater Antilles and other Pacific islands (Grehan 2001, Heads & Grehan, in press). It is clear that the only sand spider of the islands, *S. utriformis*, is of South American origins, as its closest relative lives in the coast of Peru (Fig. 2). The estimated age of split of these two species (1.2– 22 Myr, 95% highest posterior density interval, median 9.7) exceeds the age of the oldest emerged islands. Accordingly, a biogeographic model allowing for dispersal to the islands in the last 15 Myr is strongly favored in a relation to a model allowing dispersal only in the last 3.5 Myr (Fig. 6). In most trees the *S. utriformis–S. peruensis* split is older than 3.5 Myr; thus the second scenario requires that *S. utriformis* had originated in coastal Peru through in-situ speciation before 3.5 Myr, dispersed to the islands after their formation, and then went extinct in the continent (Online Supplementary Figure S13). In contrast, the first scenario only requires one dispersal event from the ancestor of the pair of species to the islands, followed by a (costless) vicariant event. Thus, our data fit better a model in which *Sicarius* reached the Galapagos when the archipelago was formed by currently drowned paleo-islands.

Interestingly, the oldest paleo-island reported by Christie et al. (1992) is only ∼530 km distant from the continent, while the current islands lie at ∼940 km away from the continent. Thus, the paleo-islands were closer to the continent, which could help explaining how a group with poor dispersal capabilities reached a volcanic archipelago. This scenario is similar to the metapopulation vicariance model that has been proposed to explain the presence of ancient, poorly dispersing groups in recent volcanic archipelagos, particularly in the Galapagos (Heads 2018, Heads & Grehan, in press). It could also explain other instances of clades whose estimated ages are older than the islands they live in, such as sheet-web weavers in the Juan Fernández Islands (Arnedo & Hormiga, in press).

Is it possible that there were even older paleo-islands in the Galapagos? Christie et al. (1992) suggested that it is very possible that the history of the archipelago could be as old as that of the volcanic hotspot, spanning 80–90 million years, and thus other uncharted paleo-islands might exist. Further evidence for this hypothesis has been recently reviewed by Heads & Grehan (in press). To account for this possibility, we compared a model where dispersal to the Galapagos was only possible in last 15 Myr to an unconstrained model where dispersal could happen at any moment—thus, considering the possibility of even older paleo-islands. The data never provides definite support to the 15-Myr model, and in those trees where the split between *S. utriformis* and *S. peruensis* is older than 15 Myr, it strongly supports the unconstrained model (Fig. 5). In addition, the inferred age of dispersal to Galapagos in the unconstrained model has some probability of being older than 15 Myr (Fig. 4). Thus, our data is unable to reject dispersal to the Galapagos before the age of the oldest recorded paleo-islands. This is in line with the opinion that the archipelago must be very old and uncharted paleo-islands exist (Christie et al. 1992).

Interestingly, we found a correlation between the support of alternative models and the ages of split (Fig. 5). There is a positive correlation between age of split and AIC weight of the unconstrained model, stronger in the models without founder-event speciation (DEC, r = 0.86; DIVA-like, r = 0.83; DIVA-like + J, r = 0.63, BAYAREA-like + j, r = 0.50). This is probably because these models favor anagenetic events, and thus the amount of time passed is correlated with opportunity for dispersal. This means that even if the split is younger than 15 Myr, but very close to the limit between time slices, the window of opportunity for a dispersal is very narrow, and thus even in these cases the unconstrained model is favored.

### Ancient connections among Neotropical dry forests

Our data indicates that the Caatinga clade reached this area as a result of a single dispersal. Such dispersal most likely originated from northern dry forests in the Caribbean and Mesoamerica, or from Andean dry forests. Interestingly, this second possibility only appears as a likely candidate when topological uncertainty is taken into account (Table 1). Such connections seem to support the hypothesis by Pennington et al. (2000) that some areas of present-day Amazonia have been replaced by xeric vegetation in the past. They compiled detailed evidence that compellingly indicates dry conditions in the region during the Quaternary. Phylogenetic patterns of other organisms, such as birds, support the hypothesis of recent connections among Neotropical dry forests (e.g. Corbett et al. 2020). Our data, on the other hand, indicates that dispersal of sand spiders to the Caatinga happened as early as in the Oligocene (Fig. 7), with no recent dispersals to other areas. Other groups of organisms display similarly ancient histories in this biome (e.g. geckos; Werneck et al. 2012). Thus, it is not unlikely that currently disjunct Neotropical dry areas might have gone through many periods of connections over the last 30 million years, and that groups with different dispersing capabilities could have responded idiosyncratically to such connections.

### Script availability

The script for replicating the analyses is available as Supplementary Material S1, or from GitHub (https://github.com/ivanlfm/BGB_BSM_multiple_trees). The code is thoroughly commented and documented, with explanations for each of the options. It has been successfully tested with two additional datasets, one of them including fossil tips, and thus we expect it to be easily adaptable to other researchers’ needs. These datasets varied between ∼30 and ∼100 taxa, ∼10 areas (with maximum range size set to ∼3) and runs included 2–3 free parameters. In each case, analyses run over 100 posterior trees were usually completed between 8 and 12 hours on a standard personal computer (Intel ® i5-5200U 2.20 GHz with 4 GB of RAM). Thus, we expect that the analyses outlined above are not too demanding computationally.

### Concluding remarks

1. Inferences of transitions among areas and age of biogeographic events using biogeographic stochastic maps benefit from running the analysis over a sample of trees, instead of a single, target tree, since they incorporate uncertainty in topology and age estimates that might be relevant to the questions at hand. We provide a broadly customizable R script to run such analyses.
2. Including phylogenetic uncertainty is especially important in models including a founder-event speciation parameter; ages of biogeographic events estimated under these models are tightly tied to cladogenetic events, such that using a single tree results in overconfident estimates that disregard the uncertainty in age estimates present in trees from the posterior distribution.
3. Our data strongly suggests *Sicarius* most likely dispersed to the Galapagos Islands before the formation of the oldest emerged island. This is congruent with geological findings that indicate that seamounts along the volcanic hotspot are former paleo-islands of the archipelago that are now drowned.
4. Our data indicate *Sicarius* dispersed into the Caatinga around 30 Mya, suggesting an ancient colonization of this area. The route of dispersal is unclear due to topological uncertainty, but most likely consisted of a northern route connecting the Caatinga to the Caribbean and Mesoamerican dry forests, or of a southern route connecting the Caatinga to the Andean dry forests.

## Acknowledgments

We thank N. Matzke for his diligence and support to the BioGeoBEARS community through the online forums. Earlier versions of the text benefited from a critical revision by U. Oliveira.

## Data Availability Statement

All the data and scripts used in our analyses is available as Online Supplementary Material:

**Supplementary File S1**. R script for running and summarizing ancestral range estimates and biogeographic stochastic maps in a sample of trees. Updated versions can be found in https://github.com/ivanlfm/BGB_BSM_multiple_trees.

**Supplementary File S2**. Maximum clade credibility phylogenetic tree representing relationships among *Sicarius* and *Hexophthalma* from the analysis of Magalhaes et al. (2019).

**Supplementary File S3**. 27000 trees from the posterior distribution from the analysis of Magalhaes et al. (2019).

**Supplementary File S4**. Phylip-formatted file with distribution ranges of *Sicarius* and *Hexophthalma*.

**Supplementary File S5**. R script for performing biogeographic model selection in our dataset.

**Supplementary Files S6–S8**. Inputs for performing the time-stratified analysis in BioGeoBEARS allowing occupation of Galapagos only in the last 15 Myr or 3.5 Myr.

**Supplementary Files S9–S12**. Supplementary table (S9) and figures (S10–S13).

## References

Akaike H. 1973. Information theory and an extension of the maximum likelihood principle. In B. N. Petrov & F. Caski (Eds.), Proceedings of the Second International Symposium on Information Theory (pp. 267–281). Budapest: Akademiai Kiado.

Arnedo M.A., Hormiga G. In press. Repeated colonization, adaptive radiation and convergent evolution in the sheet-weaving spiders (Linyphiidae) of the south Pacific Archipelago of Juan Fernandez. Cladistics.

Baker C.M., Boyer S.L., Giribet G. 2020. A well-resolved transcriptomic phylogeny of the mite harvestman family Pettalidae (Arachnida, Opiliones, Cyphophthalmi) reveals signatures of Gondwanan vicariance. J. Biogeogr. 47: 1345–1361.

Ballesteros J.A., Sharma P.P. 2019. A critical appraisal of the placement of Xiphosura (Chelicerata) with account of known sources of phylogenetic error. Syst. Biol. 68:896–917.

Binford G.J., Callahan M.S., Bodner M.R., Rynerson M.R., Núñez P.B., Ellison C.E., Duncan R.P. 2008. Phylogenetic relationships of Loxosceles and Sicarius spiders are consistent with Western Gondwanan vicariance. Mol. Phylogenet. Evol. 49:538–53.

Bremer K. 1992. Ancestral areas: A cladistic reinterpretation of the center of origin concept. Syst. Biol. 41:436–445.

Bromham L. 2019. Six impossible things before breakfast: assumptions, models, and belief in molecular dating. Trends Ecol. Evol. 34:474–486.

Cala-Riquelme F., Gutiérrez-Estrada M., Florez-Daza A.E., Agnarsson I. 2017. A new six-eyed sand spider Sicarius Walckenaer, 1847 (Araneae: Haplogynae: Sicariidae) from Colombia, with information on its natural history. Arachnology. 17:176–182.

Ceccarelli F.S., Koch N.M., Soto E.M., Barone M.L., Arnedo M.A., Ramírez M.J. 2019. The grass was greener: repeated Evolution of specialized morphologies and habitat shifts in ghost spiders following grassland expansion in South America. Syst. Biol. 68:63–77.

Christie D.M., Duncan R.A., McBirney A.R., Richards M.A., White W.M., Harpp K.S., Fox C.G. 1992. Drowned islands downstream from the Galapagos hotspot imply extended speciation times. Nature. 355:246–248.

Corbett E.C., Bravo G.A., Schunck F., Naka L.N., Silveira L.F., Edwards S. V. 2020. Evidence for the Pleistocene Arc Hypothesis from genome-wide SNPs in a Neotropical dry forest specialist, the Rufous-fronted Thornbird (Furnariidae: Phacellodomus rufifrons). Mol. Ecol. 29:4457–4472.

Drummond A.J., Ho S.Y.W., Phillips M.J., Rambaut A. 2006. Relaxed phylogenetics and dating with confidence. PLoS Biol. 4:699–710.

DRYFLOR. 2016. Plant diversity patterns in neotropical dry forests and their conservation implications. Science 353:1383–1388.

Dupin J., Matzke N.J., Särkinen T., Knapp S., Olmstead R.G., Bohs L., Smith S.D. 2017. Bayesian estimation of the global biogeographical history of the Solanaceae. J. Biogeogr. 44:887–899.

Echeverría-Londoño S., Enquist B.J., Neves D.M., Violle C., Boyle B., Kraft N.J.B., Maitner B.S., McGill B., Peet R.K., Sandel B., Smith S.A., Svenning J.-C., Wiser S.K., Kerkhoff A.J. 2018. Plant functional diversity and the biogeography of biomes in North and South America. Front. Ecol. Evol. 6:1–12.

Grehan J. 2001. Biogeography and evolution of the Galapagos: integration of the biological and geological evidence. Biol. J. Linn. Soc. 74:267–287.

Heads M. 2018. Metapopulation vicariance explains old endemics on young volcanic islands. Cladistics. 34:292–311.

Heads M., Grehan J.R. In press. The Galápagos Islands: biogeographic patterns and geology. Biol. Rev.

Ho, SYW, Duchêne S. 2014. Molecular-clock methods for estimating evolutionary rates and timescales. Mol. Ecol. 23: 5947–5965.

Hortal J., De Bello F., Diniz-Filho J.A.F., Lewinsohn T.M., Lobo J.M., Ladle R.J. 2015. Seven Shortfalls that Beset Large-Scale Knowledge of Biodiversity. Annu. Rev. Ecol. Evol. Syst. 46:523–549.

Huelsenbeck J.P., Rannala B., Masly J.P. 2000. Accommodating phylogenetic uncertainty in evolutionary studies. Science 288:2349–2350.

Huelsenbeck J.P., Imennov N.S. 2002. Geographic origin of human mitochondrial DNA: Accommodating phylogenetic uncertainty and model comparison. Syst. Biol. 51:155–165.

Lotz L.N. 2018. An update on the spider genus Hexophthalma (Araneae: Sicariidae) in the Afrotropical region, with descriptions of new species. Eur. J. Taxon. 2018:475–494.

Lüdecke D. 2018. sjmisc: data and variable transformation functions. J. Open Source Softw. 3: 754.

Magalhaes I.L.F., Oliveira U., Santos F.R., Vidigal T.H.D.A., Brescovit A.D., Santos A.J. 2014. Strong spatial structure, Pliocene diversification and cryptic diversity in the Neotropical dry forest spider Sicarius cariri. Mol. Ecol. 23:5323–5336.

Magalhaes I.L.F., Brescovit A.D., Santos A.J. 2017. Phylogeny of Sicariidae spiders (Araneae: Haplogynae), with a monograph on Neotropical Sicarius. Zool. J. Linn. Soc. 179:767–864.

Magalhaes I.L.F., Neves D.M., Santos F.R., Vidigal T.H.D.A., Brescovit A.D., Santos A.J. 2019. Phylogeny of Neotropical Sicarius sand spiders suggests frequent transitions from deserts to dry forests despite antique, broad-scale niche conservatism. Mol. Phylogenet. Evol. 140:106569.

Matzke NJ. 2013a. BioGeoBEARS: BioGeography with Bayesian (and likelihood) evolutionary analysis in R Scripts. Available from: https://github.com/nmatzke/BioGeoBEARS (last accessed October 13, 2020).

Matzke N.J. 2013b. Probabilistic historical biogeography: new models for founder-event speciation, imperfect detection, and fossils allow improved accuracy and model-testing. Front. Biogeogr. 5.

Matzke N.J. 2014. Model selection in historical biogeography reveals that founder-event speciation is a crucial process in island clades. Syst. Biol. 63:951–970.

Matzke N.J. 2021. Statistical comparison of DEC and DEC+J is identical to comparison of two ClaSSE submodels, and is therefore valid. OSF Preprint, April 27.1–40.

Nash J.C. 2014. On best practice optimization methods in R. J. Stat. Softw. 60:1–14.

Nylander J.A.A., Olsson U., Alström P., Sanmartín I. 2008. Accounting for phylogenetic uncertainty in biogeography: a Bayesian approach to dispersal-vicariance analysis of the thrushes Aves: Turdus). Syst. Biol. 57: 257–268.

Paradis E, Schliep K. 2019. ape 5.0: an environment for modern phylogenetics and evolutionary analyses in R. Bioinformatics 35: 526–528.

Parham J.F., Donoghue P.C.J., Bell C.J., Calway T.D., Head J.J., Holroyd P.A., Inoue J.G., Irmis R.B., Joyce W.G., Ksepka D.T., Patané J.S.L., Smith N.D., Tarver J.E., Van Tuinen M., Yang Z., Angielczyk K.D., Greenwood J.M., Hipsley C.A., Jacobs L., Makovicky P.J., Müller J., Smith K.T., Theodor J.M., Warnock R.C.M., Benton M.J. 2012. Best practices for justifying fossil calibrations. Syst. Biol. 61:346–359.

Pennington R.T., Prado D.E., Pendry C. a. 2000. Neotropical seasonally dry forests and Quaternary vegetation changes. J. Biogeogr. 27:261–273.

Prado D.E., Gibbs P.E. 1993. Patterns of species distributions in the dry seasonal forests of South America. Ann. Missouri Bot. Gard. 80:902–927.

R Core Team (2020). R: A language and environment for statistical computing. R Foundation for Statistical Computing, Vienna, Austria.

Rabosky D.L. 2014. Automatic detection of key innovations, rate shifts, and diversity-dependence on phylogenetic trees. PLoS One. 9: e89543.

Ree R.H., Sanmartín I. 2018. Conceptual and statistical problems with the DEC+J model of founder-event speciation and its comparison with DEC via model selection. J. Biogeogr. 45:741–749.

Ree R.H., Smith S.A. 2008. Maximum likelihood inference of geographic range evolution by dispersal, local extinction, and cladogenesis. Syst. Biol. 57:4–14.

Ree R.H., Moore B.R., Webb C.O., Donoghue M.J. 2005. A likelihood framework for inferring the evolution of geographic range on phylogenetic trees. Evolution. 59:2299–2311.

Renner S.S. 2016. Available data point to a 4-km-high Tibetan Plateau by 40 Ma, but 100 molecular-clock papers have linked supposed recent uplift to young node ages. J. Biogeogr. 43:1479–1487.

Revell LJ. 2012. phytools: an R package for phylogenetic comparative biology (and other things). Methods in Ecol. Evol. 3: 217–223.

Ronquist F. 1997. Dispersal-vicariance analysis: A new approach to the quantification of historical biogeography. Syst. Biol. 46:195–203.

Santaquiteria A., Siqueira A.C., Duarte-Ribeiro E., Carnevale G., White W., Pogonoski J., Baldwin C.C., Ortí G., Arcila D., Betancur-R. R. In press. Phylogenomics and historical biogeography of seahorses, dragonets, goatfishes, and allies (Teleostei: Syngnatharia): assessing the factors driving uncertainty in biogeographic inferences. Syst. Biol.

Suh A. 2016. The phylogenomic forest of bird trees contains a hard polytomy at the root of Neoaves. Zool. Scr. 45:50–62.

Wagenmakers E.J., Farrell S. 2004. AIC model selection using Akaike weights. Psychon. Bull. Rev. 11:192–196.

Warnock R.C.M., Parham J.F., Donoghue P.C.J., Joyce W.G., Lyson T.R. 2015. Calibration uncertainty in molecular dating analyses: there is no substitute for the prior evaluation of time priors. Proc. R. Soc. B Biol. Sci. 282:20141013.

Werneck F.P., Gamble T., Colli G.R., Rodrigues M.T., Sites J.W. 2012. Deep diversification and long-term persistence in the south american “dry diagonal”: integrating continent-wide phylogeography and distribution modeling of geckos. Evolution 66:3014–3034.

Werner R., Hoernle K., Van Den Bogaard P., Ranero C., Von Huene R., Korich D. 1999. Drowned 14-m.y.-old Galapagos archipelago off the coast of Costa Rica: Implications for tectonic and evolutionary models. Geology. 27:499–502.

White W.M., McBirney A.R., Duncan R.A. 1993. Petrology and geochemistry of the Galápagos Islands: portrait of a pathological mantle plume. J. Geophys. Res. 98:533–563.

Yu Y., Harris A.J., He X. 2010. S-DIVA (Statistical Dispersal-Vicariance Analysis): a tool for inferring biogeographic histories. Mol. Phylogenet. Evol. 56:848–850.

Yu Y., Harris A.J., Blair C., He X. 2015. RASP (Reconstruct Ancestral State in Phylogenies): a tool for historical biogeography. Mol. Phylogenet. Evol. 87:46–49.

