## Supplementary material for "Incorporating topological and age uncertainty into event-based biogeography supports paleo-islands in Galapagos and ancient connections among Neotropical dry forests": Online Supplementary

**Supplementary File S1.** R script for running and summarizing ancestral range estimates and biogeographic stochastic maps in a sample of trees.

**Supplementary File S2.** Maximum clade credibility phylogenetic tree representing relationships among *Sicarius* and *Hexophthalma* from the analysis of Magalhaes et al. (2019).

**Supplementary File S3**. 27000 trees from the posterior distribution from the analysis of Magalhaes et al. (2019).

**Supplementary File S4.** Phylip-formatted file with distribution ranges of *Sicarius* and *Hexophthalma*.

**Supplementary File S5.** R script for performing biogeographic model selection in our dataset.

**Supplementary Files S6–S8.** Inputs for performing the time-stratified analysis in BioGeoBEARS allowing occupation of Galapagos only in the last 15 Myr or 3.5 Myr.

**Supplementary Files S9–S13.** Supplementary tables (S9) and figures (S10–S13).

**Supplementary table S9**. Parameters estimated under different biogeographic models using the maximum clade credibility tree. k = number of parameters, d = dispersal, e = extinction, j = founder-event speciation.

| Biogeographic model | LnL | k | AICc | AICw | d | e | j |
|---|---|---|---|---|---|---|---|
| **DEC** | -55.83073 | 2 | 116.2328891 | 1.87E-05 | 0.001220105 | 0.002343765 | 0 |
| **DIVA-like** | -51.29909 | 2 | **107.1696254** | 0.001738148 | 0.001283512 | 1.00E-12 | 0 |
| **BAYAREA-like** | -65.06277 | 2 | 134.6969844 | 1.83E-09 | 0.001646803 | 0.022727776 | 0 |
| **DEC+J** | -44.40755 | 3 | 96.01511684 | 0.459470468 | 0.000265633 | 1.00E-12 | 0.0286 |
| **DIVA-like +J** | -44.24971 | 3 | **95.69943459** | 0.538030793 | 0.000399307 | 1.00E-12 | 0.0187 |
| **BAYAREA-like +J** | -50.83619 | 3 | 108.8723967 | 0.000741882 | 0.000285032 | 0.006871115 | 0.0269 |

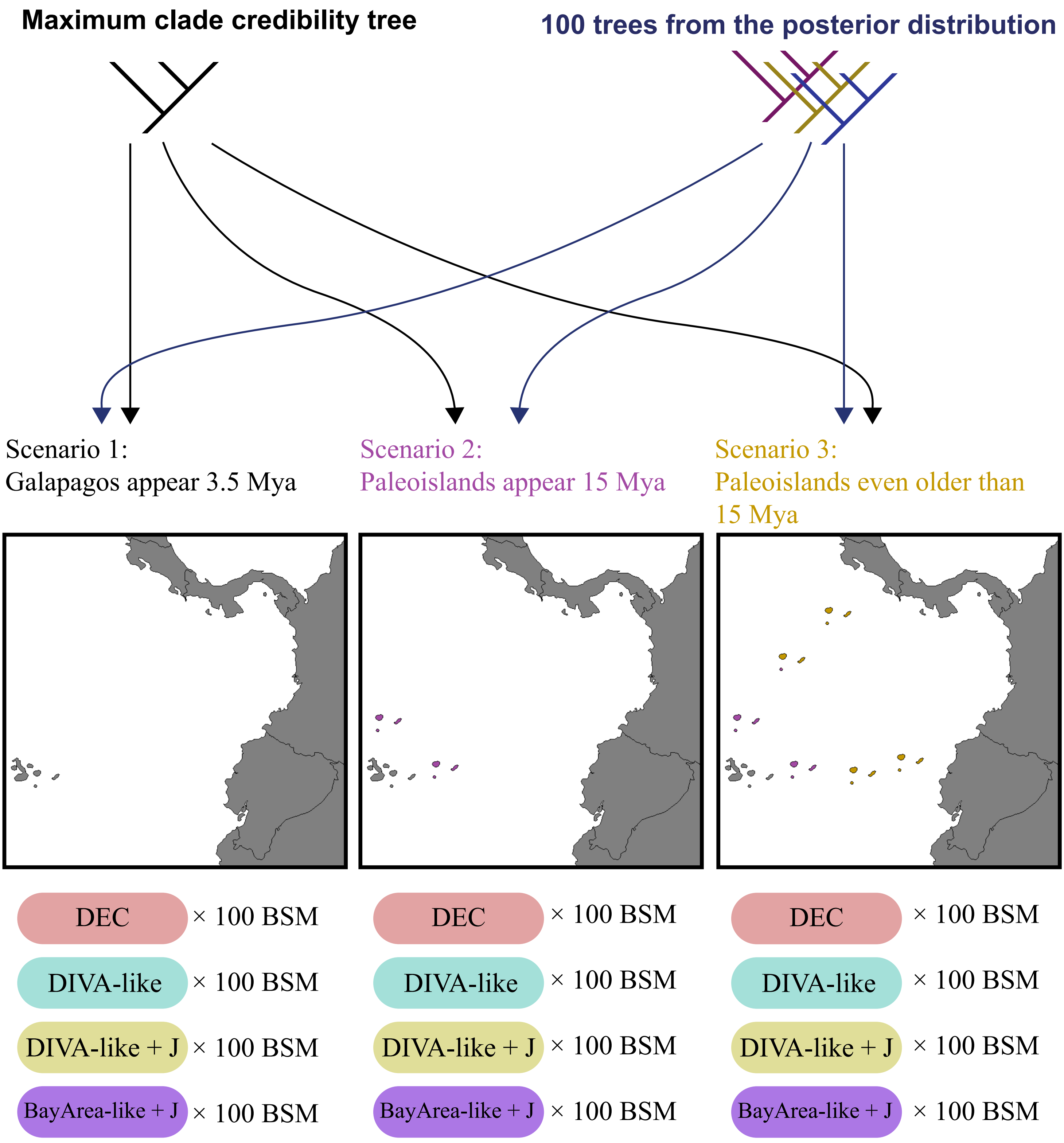


**Supplementary Figure S10.** Experimental design to investigate the effects of topology and model choice when comparing different biogeographic scenarios. We ran analysis using either the consensus tree or 100 trees taken from the posterior distribution (top row). A hundred biogeographic stochastic maps (BSM) where run for ancestral ranges estimated under four different models (bottom row). These estimates were done under three different time-stratification scenarios (middle row: Galapagos (1) only available after 3.5 Myr, (2) only available after 15 Myr, and (3) always available for occupation).

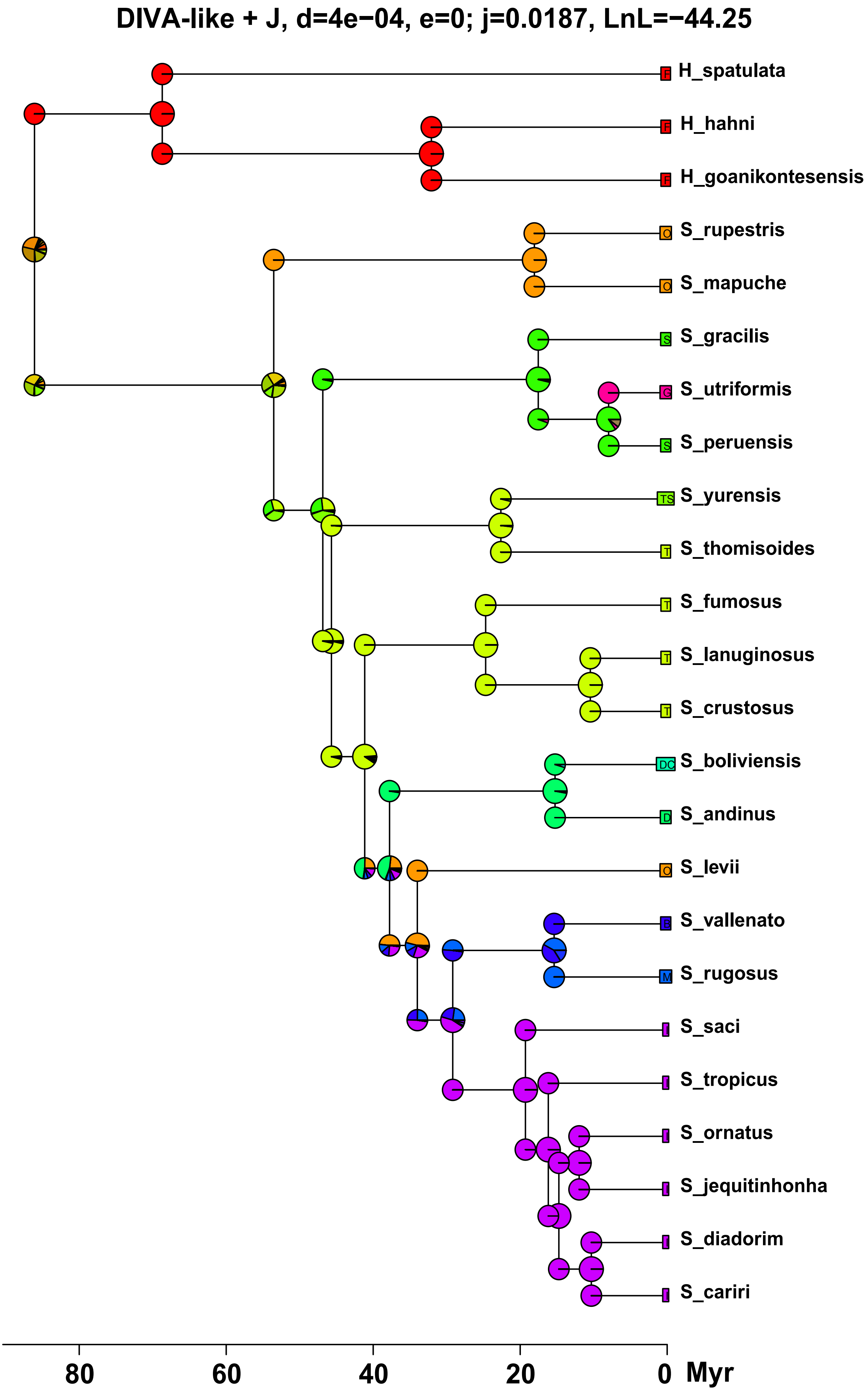


**Supplementary figure S11.** Ancestral range estimates under DIVA-like+J using the maximum clade credibility tree.

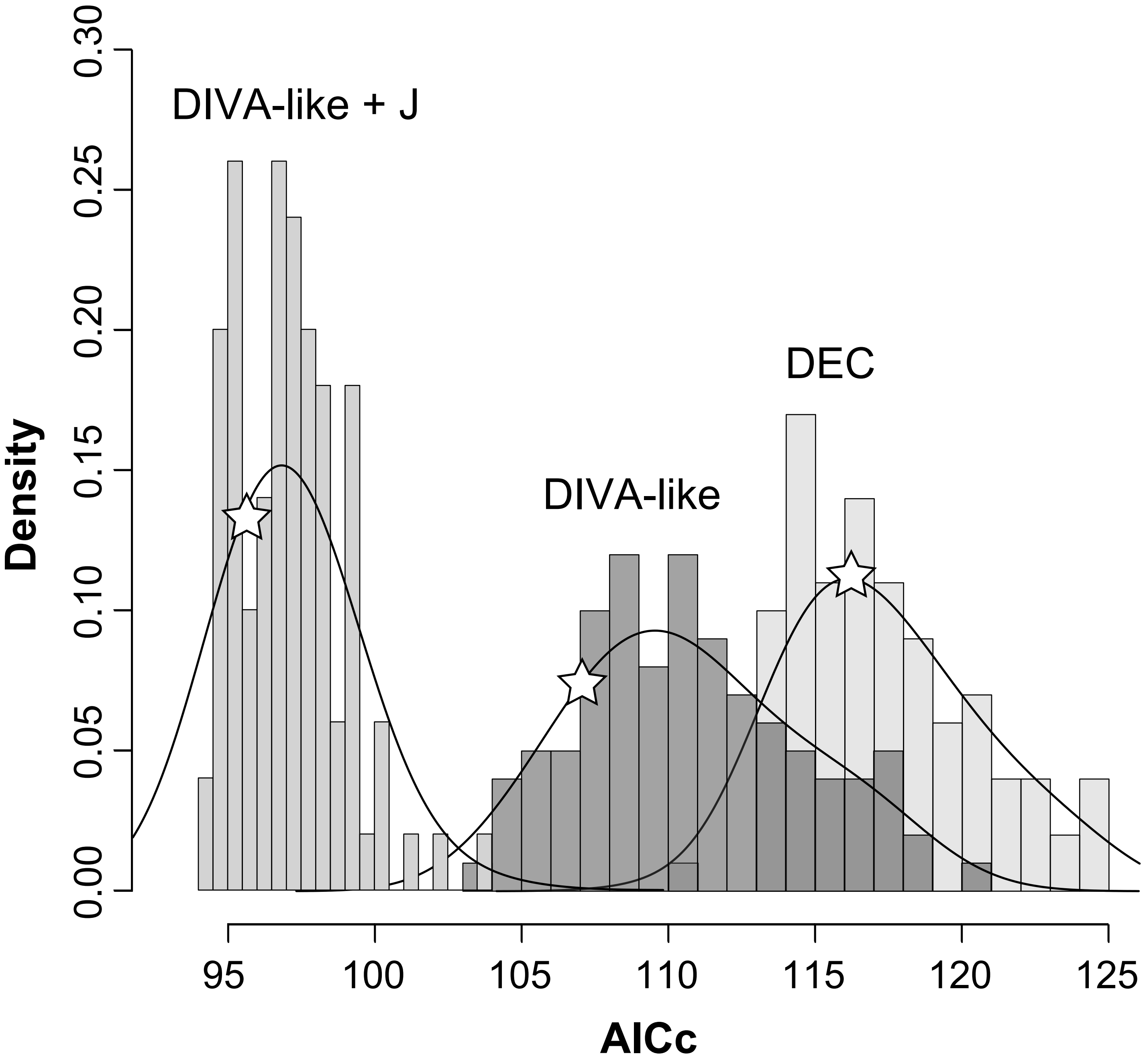


**Supplementary figure S12.** Frequency histogram of AICc values of different biogeographic models (DIVA-like+J, DIVA-like, and DEC) for each of the 100 trees sampled from the posterior distribution. The position of the MCC tree is denoted with a star.

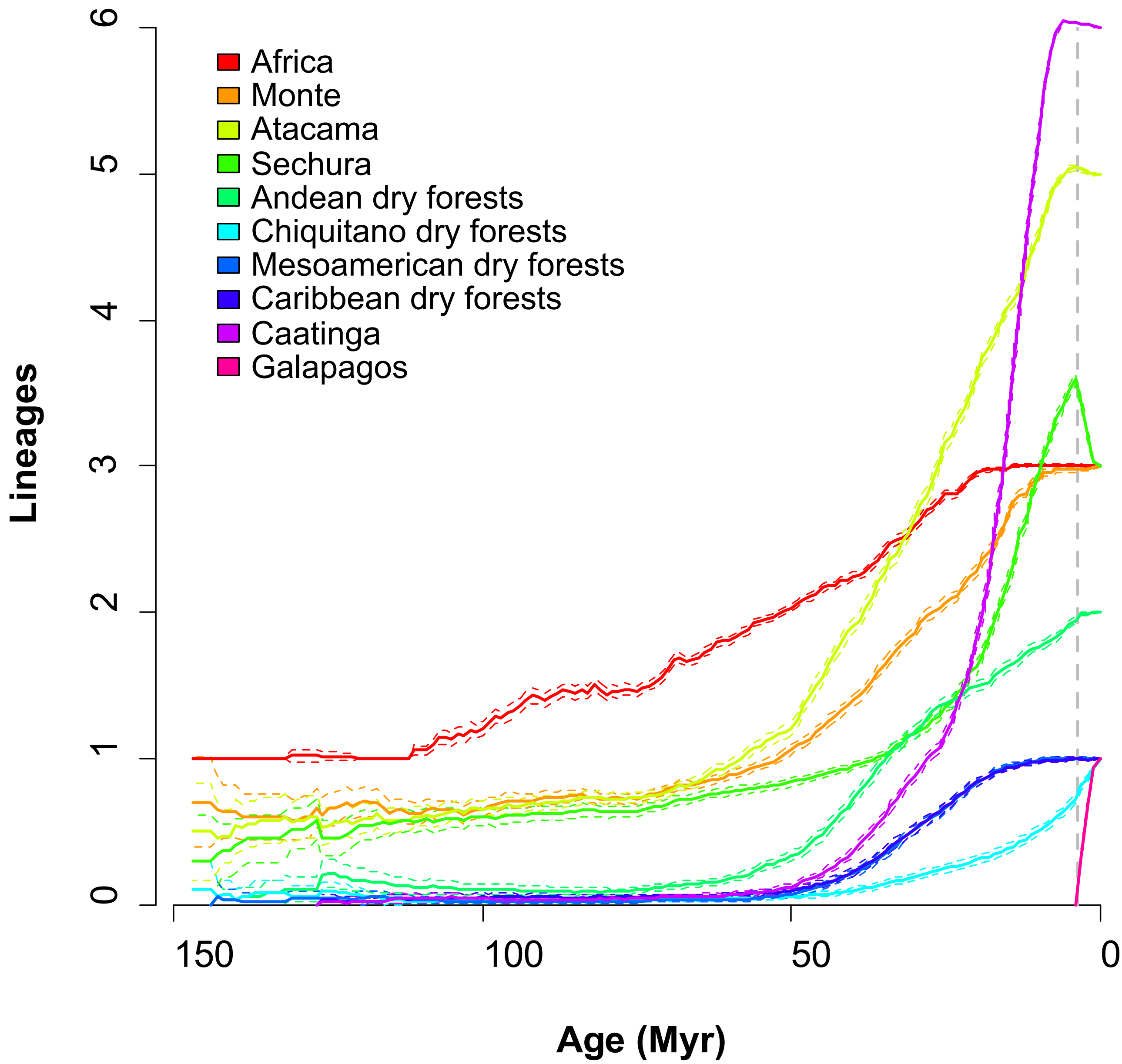


**Supplementary figure S13.** Lineages through time per area under DIVA-like, averaged over 100 trees and modeled under a time-stratified scenario where occupation of Galapagos is only possible in the last 3.5 Myr (dashed line). Note the decrease in the diversity in the Sechura desert (S). In most trees the split between *S. utriformis* (from Galapagos) and *S. peruensis* (from Sechura) is older than 3.5 Myr, causing the program to infer that *S. utriformis* originated in the Sechura desert, dispersed to Galapagos, and was later extinct from the former area. Because this requires an extra extinction event, this time-stratified scenario where occupation of Galapagos is only allowed in the last 3.5 Myr is rejected in favor of more permissive scenarios.
